# Polycomb group proteins protect latent Kaposi sarcoma-associated herpesvirus from episome clearance and HUSH-dependent chromatin silencing

**DOI:** 10.1101/2025.02.10.637455

**Authors:** Simon Weissmann, Alexis Robitaille, Thomas Günther, Marion Ziegler, Adam Grundhoff

## Abstract

Kaposi sarcoma-associated herpesvirus (KSHV) is the causative agent of a number of human neoplasms. KSHV causes lifelong chronic infections by establishing persistent reservoirs of latently infected cells in which the viral double-stranded DNA genome is present in the form of an extrachromosomal episome. Polycomb group protein (PcG) mediated gene repression is detected early after entry of the CpG-rich and epigenetically naïve viral DNA into the host nucleus. Here we show that the establishment and maintenance of KSHV latency is strongly influenced by Polycomb Repressive Complexes 1 and 2 (PRC1, and PRC2, respectively). Using a series of genetic deletion models, we dissect the functional interdependence of PRC1 and PRC2 on KSHV and identify KSHV episomes as a prime target for independent as well as collaborative *de novo* PRC recruitment. Loss of PRC function leads to a severely altered chromatin state on KSHV and we show that PRC mediated gene repression is not only preventing transcription of viral genomes but also protects KSHV episomes from detection by the human silencing hub (HUSH). Furthermore, we demonstrate that PRC complexes play an important role in ensuring episomal maintenance in dividing cells. Taken together, our results show that absence of PRC complexes creates a metastable state of KSHV latency and suggest early chromatinization as an integral step for KSHV latency establishment.

## 1. Introduction

The gammaherpesvirus KSHV is the etiological agent of Kaposi sarcoma (KS) and a range of B cell malignancies, including Primary effusion lymphoma (PEL) and multicentric Castleman’s disease (MCD) (Dittmer & Damania, 2019; Cesarman *et al*, 1995). Representing a member of the Herpesvirus family, KSHV encodes its genome as double stranded DNA, which persists as a circular episome in the nucleus of the host cell. As KSHV infects dividing cells, it ensures persistence of latent episomes *via* the viral protein LANA (Latency Associated Nuclear Antigen). The multifunctional protein LANA facilitates both recruitment of the host replication machinery for licensed DNA replication during S-Phase, but also tethers the replicated viral DNA to mitotic host chromosomes (Barbera *et al*, 2006; Juillard *et al*, 2016). Thereby KSHV causes lifelong chronic infections and establishes reservoirs of latently infected cells, contributing to tumor development (Broussard & Damania, 2020). KSHV can revert from its latent state and enter the lytic phase by expressing viral ORF50 (RTA), triggering a coordinated cascade of viral gene expression leading to extensive viral genome amplification, assembly of viral particles resulting in production of viral progeny. Chromatin modifications and chromatin associated regulatory proteins strongly impact the establishment of latency as well as lytic reactivation of herpes viruses. However, packaged viral genomes within virions are neither DNA methylated nor associated with histones. Consequently, nucleosomes and histone modifications have to be established *de novo* upon entry of the naked viral DNA into the host nucleus. The timely establishment of histone modifications ensures transcriptional repression of the majority of the KSHV genome, while only genes involved in latency maintenance continue to be expressed.

Polycomb Repressive Complexes (PRCs) play a significant role in the chromatinization and latency establishment of KSHV. Early after infection and in established tumors, the majority of the KSHV genome is decorated by polycomb-mediated histone marks (Günther & Grundhoff, 2010; Toth *et al*, 2010, 2013; Günther *et al*, 2014; Sun *et al*, 2017; Günther *et al*, 2019). High levels of PRC mediated histone marks have been proposed as a viral strategy to repress lytic gene expression during latency (Günther & Grundhoff, 2010; Toth *et al*, 2010, 2013).

Polycomb group proteins (PcG) are evolutionary conserved regulators of transcription, associated with chromatin and involved in the modification of gene expression during the development of higher eukaryotes (Schuettengruber *et al*, 2017; Bracken *et al*, 2006; Margueron *et al*, 2008; Piunti & Shilatifard, 2021). They exert their gene-repressive function through two multimeric complexes, Polycomb Repressive Complex 1 (PRC1) and Polycomb Repressive Complex 2 (PRC2). These complexes contain enzymatically active cores that, along with auxiliary proteins, form distinct subcomplexes modulating PRC activity and recruitment (Blackledge & Klose, 2021).

The catalytic core of PRC2 contains EED, SUZ12 and EZH1 or its homolog EZH2, which associate with equimolar stoichiometries and together with RBBP4 or RBBP7 can bind additional interaction partners to form the subcomplexes PRC2.1 and PRC2.2 (Glancy *et al*, 2021; Laugesen *et al*, 2019). Within the PRC2 core, the SET-domain containing methyltransferases EZH1 and EZH2 catalyze mono-, di- and tri-methylation of lysine 27 of histone H3 (H3K27me1, H3K27me2 and H3K27me3) (Cao *et al*, 2002; Ferrari *et al*, 2014; Margueron *et al*, 2008). Subcomplex variant PRC2.1 contains one polycomb-like protein (PCL1, PCL2 or PCL3) as well as either PALI1/2 or EPOP, whereas PRC2.2 contains Jumonji and (A+T)-rich interaction domain-containing protein 2 (JARID2) and adipocyte enhancer-binding protein 2 (AEBP2) (Beringer *et al*, 2016; Poepsel *et al*, 2018).

PRC1’s catalytic core contains the E3 ubiquitin ligase RING1B or its paralog RING1A (also known as RNF2 and RING1 respectively) which catalyze the mono-ubiquitination of histone H2A at lysine 119 (H2AK119Ub). RNF2 and RING1 further associate with one of six Polycomb group RING finger (PCGF) proteins (Blackledge & Klose, 2021; Gao *et al*, 2012). PRC1 variants are categorized as “canonical” (cPRC1) or “non-canonical” (PRC1.1 - PRC1.6) complexes, with cPRC1 defined by the presence of a chromodomain-containing CBX subunit, while all non-canonical PRC1 complexes contain RYBP or YAF2 (Piunti & Shilatifard, 2021).

The mechanism of PRC recruitment in mammalian cell systems has long been poorly defined. While PRC targeting in *D.melanogaster* is facilitated by sequence specific polycomb recruitment elements (PRE), retention and enrichment of polycomb complexes in mammals is thought to involve CpG islands (CGI). Low levels of PRC1 and PRC2 mediated marks are globally dispersed on the mammalian genome, most likely originating from intrinsic weak interaction of PRC complexes to DNA (Højfeldt *et al*, 2018; Fursova *et al*, 2021). However, high levels of both marks and respective complexes are focused on unmethylated CGIs of genes lacking transcription (Riising *et al*, 2014; Wachter *et al*, 2014). Two hierarchical models of focusing PRC mediated marks have been proposed – both involving PRC subunits that increase retention time *via* CpG binding domains: The non-canonical PRC1.1 complex can bind stretches of unmethylated CpGs *via* its subunit KDM2B (Wu *et al*, 2013; Farcas *et al*, 2012). PRC1.1 harbors high catalytic activity, placing H2AK119Ub which can be detected by AEBP2/JARID2 containing PRC2.2, thereby enhancing H3K27me3 deposition (Kasinath *et al*, 2021). Conversely, the canonical PRC recruitment pathway is characterized by initial stabilization of PRC2.1 binding to CGIs via its PCL1-3 complex members, deposition of H3K27m3, which in turn can be recognized by the CBX subunits of cPRC1 triggering H2AK119Ub deposition (Rowitch & Kriegstein, 2010). Importantly, both recruitment scenarios rely on the presence of stretches of unmethylated CpG dinucleotides. We have previously shown that KSHV harbors a high density of CpG motifs and KDM2B binding is found early during infection indicating that non-canonical PRC1.1 driven recruitment is a major axis of PRC function on KSHV (Günther *et al*, 2019). However, the exact interplay of PRC1 and PRC2 on KSHV are poorly understood. Furthermore, the functional consequences of complete loss of Polycomb function on KSHV episomes are unknown.

Interestingly, early after infection, the KSHV genome is not decorated by classical heterochromatin marks such as H3K9me3, which linked to transcriptional repression of foreign DNA. The absence of H3K9me3 suggests that KSHV employs unique mechanisms to evade detection by host antiviral defenses. Furthermore, the role of the HUSH complex, a mediator of transcriptional silencing of foreign DNA through H3K9me3 deposition SETDB1 (Seczynska *et al*, 2022; Seczynska & Lehner, 2023), has not been investigated in the context of KSHV infection.

In this study we aimed at elucidating the role of Polycomb repressive complexes in KSHV infection. In particular we investigated their role in mediating gene repression on KSHV, as well as their importance in establishing a chromatin state which supports latency and episomal maintenance. We present a systematic dissection of the different PRC components using combinatorial genetic deletion models, a comprehensive analysis of KSHV chromatin in the absence of PRC complexes and observe a strong dependency of PRC complexes to ensure episomal maintenance.

## 2. Materials and Methods

### 2.1 Cell culture, Transfection, Lentivirus transduction

SLK, EA.hy926 (ATCC), HEK293 and HAP1 cells (Horizon Discovery) were cultivated in DMEM High Glucose (Gibco) supplemented with 10 % FCS, 1 mM Sodium Pyruvate (Fisher Scientific) 2 mM L-Glutamine, 100 U/ml Penicillin and 100 μg/ml Streptomycin. BCBL1 cells were cultured in RPMI 1640 medium containing 10 % FCS, 100 U/ml Penicillin and 100 μg/ml Streptomycin. SLK has been discovered to be a misidentified cell line (Cellosaurus AC: CVCL_9569) and was identified as a contaminating cell line (Caki-1; Cellosaurus AC: CVCL_0234) (Stürzl *et al*, 2013)

All cell lines were regularly tested for mycoplasma contaminations using LookOut Mycoplasma PCR Detection Kit (Sigma-Aldrich, MP0035–1KT).

Transient transfection was performed via reverse transfection with LT1 transfection reagent (Mirus) for HAP1 cells and via electroporation for SLK cells (AMAXA Nucleofector II program A20 - LONZA with Ingenio® Electroporation Solution -Mirus).

Lentiviral particles were produced in Lenti-X™ 293T cells with second-generation packaging plasmids (psPAX2 and pCMV-VSV-G). SUZ12 rescue experiments were performed by cloning human SUZ12 cDNA (Addgene # 24232) into a LeGO lentiviral vector (Weber *et al*, 2008). *SUZ12*^KO^ HAP1 clones were transduced with LeGO-iGFP-blast-SUZ12-V5, selected with blasticidine (10 µg/ml, InvivoGen) and sorted for high GFP expression.

### 2.2 Generation of KO clones

All knockout cell lines were generated by transient transfection of pSpCas9(BB)-2A-GFP (PX458, addgene # 48138), using two gRNAs per target gene. Clonal cell lines, established by FACS, were screened by PCR (Phire Tissue Direct PCR, Thermo Fisher) and gene deletion was confirmed by immunoblotting. Combinatorial deletions were established from single knockout lines. gRNA primer sequences used for knockout generation are listed in **Supplementary Table S1**. Unless otherwise stated, replicate experiments of infection experiments were performed in two independent clones.

### 2.3 KSHV production and infection

Infectious wild-type KSHV supernatant was produced by inducing latently infected BCBL1 cells with TPA (Sigma-Aldrich; Cat#P8139–5MG) and sodium butyrate (Sigma-Aldrich; Cat#B5887–1G) followed by 100-fold concentration by centrifugation as described previously (Grundhoff & Ganem, 2004). rKSHV.219 was produced from the latently infected Vero cell line VK219.

An ORF50 deletion mutant (KSHV-BAC16-ΔORF50) of KSHV-BAC16 (Brulois *et al*, 2012) was generated by *en passant* mutagenesis (Tischer *et al*, 2010), where the reading frame, but not the 5’ UTR, of ORF50 was replaced with mCherry. The resulting KSHV-BAC16-ΔORF50 was analyzed by restriction enzyme analysis with multiple restriction enzymes and long read whole plasmid sequencing (Eurofins Genomics).

For reconstitution of KSHV-BAC16 and KSHV-BAC16-ΔORF50, bacmid constructs were transfected into iSLK cells. Virus production was performed as previously described (Brulois *et al*, 2012). For infection, concentrated KSHV stock solutions were diluted in EBM-2 medium (LONZA) containing 8 µg/ml polybrene, and incubated with target cells for two hours. KSHV-BAC16 infected cells were selected with hygromycine B (200 µg/ml, InvivoGen) for one week to achieve 100 % KSHV+ cultures.

### 2.4 Chromatin Immunoprecipitation (ChIP)

ChIP was performed as described previously (Günther & Grundhoff, 2010). Cells were cross-linked (1 % formaldehyde, 10 minutes), quenched with 125 mM glycine, washed twice with PBS and harvested in 1 ml buffer 1 (50 mM Hepes-KOH, 140 mM NaCl, 1 mM EDTA, 10 % glycerol, 0.5 % NP-40, 0.25 % Triton X-100) and incubated for 10 min at 4 °C while rotating. After centrifugation (1,350 x g, 5 min), nuclei were incubated with 1 ml buffer 2 (10 mM Tris-HCl, 200 mM NaCl, 1 mM EDTA, 0.5 mM EGTA) for 10 min at 4 °C while rotating. Pelleted nuclei were lysed in buffer 3 (1 % SDS, 10 mM EDTA, 50 mM Tris-HCl). Chromatin was sonicated (fragment size 200-500 bp) using a Bioruptor™ (Diagenode). After addition of Triton X-100 (1 % final concentration) cell debris was pelleted (20,000 x g, 4°C) and chromatin containing supernatant was collected. Chromatin of 1×10^6^ cells was diluted 1:10 in dilution buffer (0.01 % SDS, 1.1 % Triton X-100, 1.2 mM EDTA, 16.7 mM Tris-HCl, 167 mM NaCl) and incubated with respective antibodies overnight. 50 μl BSA-blocked Protein A/G Magnetic Beads (Pierce™) were added to precipitate the chromatin-immunocomplexes and incubated for 3 hr at 4°C. Beads were washed once with 1ml of the following buffers: low-salt buffer (0.1 % SDS, 1 % Triton X-100, 2 mM EDTA, 20 mM Tris-HCl, 150 mM NaCl); high-salt buffer (0.1 % SDS, 1 % Triton X-100, 2 mM EDTA, 20 mM Tris-HCl, 500 mM NaCl); LiCl-wash buffer (0.25 M LiCl, 1 % Nonidet P-40, 1 % Na-deoxycholate, 1 mM EDTA, 10 mM Tris-HCl) and TE-wash buffer. Chromatin was eluted and decrosslinked from the beads by incubation in 120 μl SDS containing elution-buffer (50 mM Tris-HCl pH 8.0, 10 mM EDTA, 1 % SDS) containing 200 mM NaCl at 65 °C overnight. Chromatin containing supernatant was separated from the beads by a magnetic rack. DNA was purified using a DNA Clean & Concentrator kit (Zymo Research). For ChIP-qPCR, DNA was analyzed using the Rotor-Gene real-time PCR cycler (QIAGEN) with SensiMix™ SYBR® Hi-ROX (BioCat). For ChIP-seq, 1–2 ng of ChIP DNA was used for library preparation, using the NEBNext Ultra II DNA Library prep Kit (E7370; NEB). Libraries were sequenced using Illumina NextSeq 500 sequencer 75 bp Single End or Illumina NextSeq 2000 sequencer 50 bp Single End. Primers and antibodies are listed in **Supplementary Table S1.**

### 2.5 RNA Quantification and RNA-seq

RNA was isolated using TRIzol™ Reagent (Invitrogen) according to manufacturer’s recommendation. For cDNA, 2 µg RNA was treated with DNAse I (Life Technologies) and RNA was reverse transcribed using SuperScript™ Reverse Transcriptase (Invitrogen) and random Hexamers (Eurofins Genomics). cDNA was quantified using the Rotor-Gene real-time PCR cycler (QIAGEN) with SensiMix™ SYBR® Hi-ROX (BioCat). RNA samples without Reverse Transcriptase treatment served as negative controls. Gene expression was normalized to two housekeeping genes (*RPLP0*, *PLK1*). Gene expression of KSHV ORFs was further normalized to episomal content. For RNA-seq, quality of extracted total RNA was verified prior to library preparation on a 2100 Bioanalyzer (Agilent). For RNAseq libraries prepared using the CORALL RNA-Seq V2 Library Prep Kits (Lexogen) and sequenced on an Illumina NextSeq 2000 sequencer 35 bp Paired End.

### 2.6 Immunoblotting

For whole-cell extracts, cells were lysed in high salt buffer (300 mM NaCl, 50 mM Tris-HCl pH7.5, 0.5 % Triton X-100, 0.1 % SDS, 1 mM EDTA, 1 mM DTT, 1× cOmplete™ (Roche)). Equal amounts of whole-cell extracts were analyzed by standard SDS-PAGE/immunoblotting. Primary antibodies and dilutions are listed in **Supplementary Table S1**.

### 2.7 Immunofluorescence and positive cell quantification

Cells were fixed on polymer 8-well coverslips ibiTreat (#80826; ibidi) with 4 % paraformaldehyde in PBS for 10 min, permeabilized with 2 % Triton X-100 in PBS for 30 min and blocked with 3 % bovine serum albumin in PBS for 30 min. Primary antibody was diluted as listed in **Supplementary Table S1** in 3 % BSA-PBS and incubated for 2 h at room temperature and stained with secondary antibody room temperature for 1 h. After nuclear stain with Hoechst 33342 solution (Sigma Aldrich) samples were imaged at a Leica DMI6000 B microscope. To quantify fractions of positive cells, automated positive cell detection was performed in QuPath (version 0.3.0) (Bankhead *et al*, 2017). Final positive cell fractions were pooled from eight images (four images per clone of respective genotype).

### 2.8 DOX-RTA induction

KSHV cDNA of RTA/ORF50 was cloned into a doxycycline inducible piggyBac Vector (Addgene # 97421) creating pTet-ON-kRTA. HAP1 cells were transfected with pTet-ON-kRTA and transposase expressing plasmid pCMV-hyPBase (Sanger Plasmid Repository). HAP1 cells were infected with KSHV and RTA expression was induced by addition of doxycycline (1 µg/ml) 24 h post infection. Gene expression and ORF59 staining was performed 48 h post infection.

### 2.9 Episome quantification and loss experiments

Cells were infected *de novo* with KSHV, rKSHV or BAC16. Genomic DNA (gDNA) was isolated by DNeasy Blood & Tissue Kit (QIAGEN). DNA was eluted in DEPC H2O 1:1 mixed with AE buffer. Quantitative PCR (qPCR) of gDNA was performed on a Rotor-Gene real-time PCR cycler (QIAGEN) from 90 ng gDNA with SensiMix™ SYBR® Hi-ROX (BioCat) using three different primer pairs for KSHV as well as human DNA quantification. KSHV episomes were normalized to human DNA. For episomal loss experiments KSHV episome numbers were normalized to first sample (Day 2) post infection. KSHV-BAC16 infected cells were selected with hygromycine B (200 µg/ml, InvivoGen) for one week to achieve 100 % KSHV+ cultures. After removal of hygromycine B following episome quantifications were normalized to Day 0.

For pGTR8 retention assays, SLK and HAP1 cells were transduced with KSHV LANA containing lentivirus (LeGO-mCherry-kLANA) and FACS sorted for mCherry. 2 µg of pGTR8 were transfected into SLK or HAP1 cells ectopically expressing LANA and pGTR8 retention was measured by FACS by GFP positive cell fraction and normalized to first sample (Day 2 after transfection).

To determine statistical significance, slopes (k) of exponential regression were compared using two-tailed t-test when comparing two conditions. For comparing multiple slopes against one control condition, ordinary one-way ANOVA with multiple comparison correction (Dunnett) was performed.

### 2.10 Replication assay

#### BrdU incorporation assay

Asynchronously growing cultures were pulse labelled with 5-Bromo-2’-deoxyuridine (BrdU, Sigma) at 100 µM for 1h. Unlabeled cells were used as negative control. DNA was isolated using a DNeasy Blood & Tissue Kit (QIAGEN), sonicated using an NGS bioruptor (Diagenode), heat denatured (95°C, 5min) and incubated with anti-BrdU (BD Biosciences) in IP buffer (PBS with 0.0625 % Triton-X 100). Antibody bound DNA was precipitated with Protein A/G magnetic Beads (Life Technologies, blocked with 1 mg/ml BSA and 0.25 mg/ml salmon sperm DNA) and eluted from beads in elution buffer (100 mM Tris-HCl, pH 8, 1% SDS) at 65°C for 15 min. DNA was purified using a DNA Clean & Concentrator kit (Zymo Research). Purified DNA was quantified on a Rotor-Gene real-time PCR cycler (QIAGEN) with SensiMix™ SYBR® Hi-ROX (BioCat) using three different primer pairs for KSHV as well as human DNA quantification.

#### EdU incorporation assay

Asynchronously growing cultures were pulse labelled with 5-Ethynyl-2’-deoxyuridine (EdU, Sigma) at 25 µM for 1h. Unlabeled cells were used as negative control. 1×10^6^ cells were harvested and lysed in 500µl lysis buffer (50mMTris–HCl pH 7.5, 150mM NaCl, 5 mM CaCl2, 1% Triton X-100, 0.5% NP40). Chromatin containing fraction was collected by centrifugation at 500 rcf for 5 min, resuspended in 50 µl Click-iT™ Cell Reaction Buffer (Thermo Fisher, C10269) containing 10 µM Biotin-Azide (baseclick, BCFA-003) and incubated at room temperature for 25 min. Nuclei were washed once with lysis buffer and DNA was extracted using a DNeasy Blood & Tissue Kit (QIAGEN). DNA was sonicated using an NGS bioruptor (Diagenode) and biotin containing DNA was precipitated with MyOne C1 Beads (Invitrogen), washed three times with Binding and Wash buffer (BW) (5 mM Tris, pH 7.5, 0.5 mM EDTA, 1M NaCl) and eluted in elution buffer (2% SDS, 3mM Biotin in PBS) for 15min at room temperature followed by 15 min at 96 °C. DNA was purified using a DNA Clean & Concentrator kit (Zymo Research). Purified DNA was quantified on a Rotor-Gene real-time PCR cycler (QIAGEN) with SensiMix™ SYBR® Hi-ROX (BioCat) using three different primer pairs for KSHV as well as human DNA quantification.

### 2.11 Apoptosis assay

Cell fractions of apoptotic (Annexin V+) and necrotic (Annexin V+/PI+) were measured by Annexin V staining using a Pacific Blue™ Annexin V Apoptosis Detection Kit (BioLegend) and a BD LSRFortessa™ Cell Analyzer.

### 2.12 Competitive growth assay

HAP1 cells infected with KSHV-BAC16 and selected with Hygromycin were mixed with uninfected HAP1 cells expressing BFP from lentiviral vectors (LeGO-BFP) at a 9:1 ratio. Mixed cultures were passaged routinely and composition of KSHV-BAC16 (GFP+), KSHV-BAC16 cleared (GFP-) or uninfected control samples (BFP+) was monitored by FACS on a BD LSRFortessa™ Cell Analyzer.

### 2.13 Bioinformatic analysis and data availability

For ChIPseq, quality filtered single end reads were aligned to the viral reference genome of KSHV (HQ404500) and human (GRCh38) using Bowtie (Langmead *et al*, 2009) with standard settings. Quantification and analysis of histone modification enrichment on the KSHV genome was done as described in (Günther *et al*, 2019) and https://github.com/VirusGenomics/ViPeaQ. Host-positive sites were identified using peak calling with MACS2 (Zhang *et al*, 2008) for H3K27ac and EPIC2 (Stovner & Sætrom, 2019) for H3K27me3, H2AK119Ub, and H3K9me3. The 200 most significantly enriched regions were selected as host-positive regions. Host-background regions, representing general ChIP background, were generated by randomly selecting regions matching the chromosome and size distribution of the positive controls. Regions with fewer than 10 reads per kilobase (kb) in the input or known mappability biases loci (blacklisted regions from ENCODE for GRCh38) were excluded. The viral genome was divided into overlapping 10 kb windows (shifted by 5 kb) to capture broad regions. Raw reads were counted for positive, background, and viral regions using FeatureCounts (Liao *et al*, 2014) and normalized to reads per kilobase (RPK).

For each region, the ChIP-to-input RPK ratio was calculated. To enable cross-experiment comparisons, a normalization factor was applied to adjust the median RPK value of the background distribution to 1. Positive and viral distributions were normalized using the factor derived from their respective background regions. Since all distributions (positive, background, and viral) were normalized for (1) read per kilobase, (2) ChIP efficiency, and (3) the median of the background distribution, background controls are not displayed in the figures. The resulting values reflect relative enrichment of histone modification signals over the general ChIP background for each experiment. Differences in histone marks across 10 kb viral genome segments were assessed using repeated measures ANOVA and Fisher’s LSD.

For RNA-seq, quality filtered paired-end reads were aligned to the viral reference genome of KSHV (HQ404500) and human (GRCh38) using STAR (Dobin *et al*, 2013) with standard settings. For expression analysis of KSHV encoded ORFs, all ORFs annotated in HQ404500 (GenBank) were assigned as single exons and genes in GFF3 format. Aligned reads were counted using FeatureCounts (Liao *et al*, 2014) and ciral reads were normalized to episome numbers quantified by qPCR as described in Material and Methods section *Episome quantification and loss experiments*. Differential gene expression analysis of host genes (GRCh38) and KSHV (HQ404500) was performed using DESeq2 (Love *et al*, 2014). For visualization of KSHV gene expression, significant expression levels changes (P.adj < 0.05) of each KSHV ORF are plotted as heatmaps with color gradients of log2 fold change vs. WT.

The data for this study have been deposited in the European Nucleotide Archive (ENA) at EMBL-EBI under accession number PRJEB85570 (https://www.ebi.ac.uk/ena/browser/view/PRJEB85570).

## 3. Results

### 3.1 Canonical as well as non-canonical PRC recruitment determines H3K27me3 and H2AK119ub on KSHV

In order to investigate the temporal recruitment of the PRC machinery, and to identify mechanistically dependencies of PRC complex members in the context of KSHV, we decided to use cell models harboring genetic deletions for individual PRC components and investigate PRC function on KSHV in *de novo* infection models (**Figure 1A**). Previous studies have utilized RNA interference (Toth *et al*, 2013; Hopcraft *et al*, 2018), pharmacological inhibition (Toth *et al*, 2010) or overexpression of H3K27me3 specific demethylases (Günther & Grundhoff, 2010) to investigate PRC function during KSHV infection. In contrast to these approaches knockout models offer reduced confounding effects such as incomplete removal of target proteins or off-target effects. As a cellular model system, we chose HAP1 cells, which are easily genetically tractable and tolerate deletions of PRC2 (Carette *et al*, 2011; Wassef *et al*, 2019). Furthermore, they proved to be permissive for latent KSHV infection, with comparable rates of LANA positive cell fractions after *de novo* infections compared to the widely used cellular model of SLK cells (**Supplemental Figure 1E)**. We used CRISPR/Cas9 to generated stable knockout cell lines that are devoid of PRC2 (*SUZ12^KO^*) or PRC1 function (*RING1^KO^*/*RNF2^KO^*). Furthermore, we generated a cell line that harbors a triple knockout of SUZ12, RING1 and RNF2 (*RING1^KO^*/*RNF2^KO^*/*SUZ12^KO^*), where all PRC functions are abrogated (**Figure 1B**)

**Figure 1:**
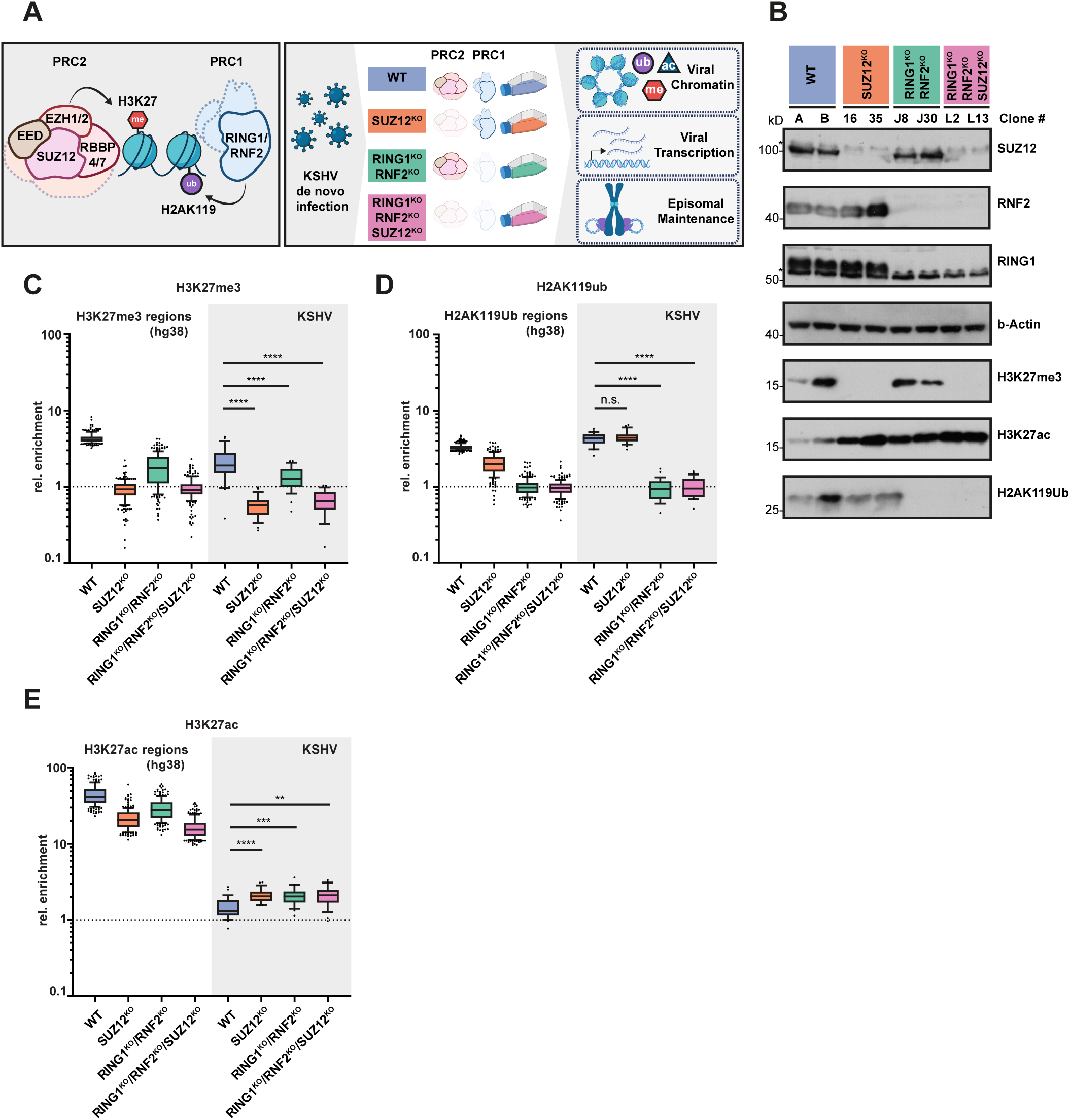
Canonical as well as non-canonical PRC recruitment is driving H3K27me3 and H2AK119ub on KSHV. **A)** Schematic overview of Polycomb function and experimental approach **B)** Immunoblotting of individual HAP1 PRC1 and PRC2 knockout clones. Asterisk symbols (*) denote unspecific bands **C-E)** Quantification of ChIP seq signal in a 10 kb sliding window over the KSHV genome (48h p.i.), normalized to negative host regions, sequencing depth and input. Significance of differences of histone marks of viral sequences segments (10kb) were calculated using repeated measure (RM) ANOVA Fisher’s LSD. P > 0.05 (n.s), P ≤ 0.05 (*), P ≤ 0.01 (**), P ≤ 0.001 (***), P ≤ 0.0001 (****)

Next, we used KSHV produced in PEL cells (BCBL1) for *de-novo* infection of our established KO cell lines, and investigated the recruitment of PRC1 and PRC2 to KSHV by detection of their respective marks, H2AK119Ub and H3K27me3, as well as the PRC2 structural component SUZ12. Using chromatin immune precipitation followed by quantitative PCR (ChIP-qPCR) or high throughput sequencing (ChIP-seq) we aimed to determine the recruitment capabilities of both cPRC1 and PRC1.1 mediated recruitment to KSHV genomes. This allowed for simultaneous measurement and comparison of PRC recruitment patterns to the human host genome.

First, we investigated H3K27me3 levels and observed the expected loss of H3K27me3 in *SUZ12^KO^* cell lines. Conversely, H3K27me3 was strongly reduced but still detectable on KSHV in *RING1^KO^*/*RNF2^KO^*(**Figure 1C, Supplemental Figure S1A, S1B**). This argues for a mechanism of PRC2 recruitment independent of PRC1 and H2AK119Ub, possibly through the stabilization of PRC2 binding to CpG islands *via* polycomb like proteins (PCL1-2, (Li *et al*, 2017)). Indeed, SUZ12 continued to localize to KSHV episomes in the complete absence of PRC1 (*RING1^KO^*/*RNF2^KO^*) (**Supplemental Figure S2A)**.

Next, we focused on determinates of H2AK119Ub establishment on KSHV. To expand our analysis to non-canonical PRC1.1 we generated HAP1 cells deficient for KDM2B (*KDM2B^KO^*). In addition, we combined the deletion of SUZ12 with a deletion of KDM2B to generate a cell line that is deficient in PRC2 as well as PRC1.1 (*KDM2B^KO^*/*SUZ12^KO^*), (**Supplemental Figure S1C, S1D**).

Loss of KDM2B had little impact on H2AK119Ub levels on KSHV (**Supplemental Figure S2A)**. Similarly, loss of SUZ12 had little consequences on the levels of H2AK119Ub on KSHV (**Figure 1D, Supplemental Figure S1A, S1B, S2A)**, which was in stark contrast to human positive control regions. This observation indicates that H2AK119Ub can be deposited by cPRC1 as well as PRC1.1. Interestingly, however, even the combined deletion of SUZ12 and KDM2B did not cause a strong reduction of H2AK119Ub on KSHV, meaning that other non-canonical PRC1 complexes, independent of KDM2B, are able to deposit H2AK119Ub on KSHV and can substitute for PRC1.1 once KDM2B is deleted (**Supplemental Figure S2A)**. Importantly, deletion of core members of both PRC1 and PRC2 complexes (*RING1^KO^*/*RNF2^KO^*/*SUZ12^KO^*) caused complete lack of H3K27me3 and H2AK119Ub on KSHV (**Figure 1C-D, Supplemental Figure S1A, S1B**).

Taken together, we show that KSHV genomes represent formidable targets for PRC recruitment, where different recruitment pathways converge to work in concert.

### 3.2 Depletion of PRC function causes hyperacetylation of H3K27 at KSHV episomes

We speculated that once PRC2 mediated deposition of H3K27 methylation is impaired this would lead to an increase of acetylation (H3K27ac). Indeed, loss of SUZ12 and consequential loss of H3K27me3 caused only minimal reduction of H2AK119Ub but drastic increase in H3K27ac (**Figure 1B, 1E)**. Similarly, loss of H2AK119Ub accompanied by reduction of H3K27me3 resulted in increased H3K27ac, although to a lesser degree than PRC2 deletion (**Figure 1E, Supplemental Figure S1A, S1B)**. In fact, increase of H3K27ac levels robustly mirrored the degree of H3K27me3 loss on KSHV. In an effort to corroborate our results obtained by knockout-cell lines, we used EPZ-6438, a highly potent small molecule inhibitor of EZH1/2, which is approved for treatment of B-cell malignancies (Knutson *et al*, 2013) (**Supplemental Figure S2C**). Similar to genetic ablation of PRC complex members, treatment with EPZ-6438 during *de novo* infection with KSHV caused loss of H3K27me3 and increase of H3K27ac **(Supplemental Figure S2D)**. Reconstitution of SUZ12 in a *SUZ12^KO^*cell line (**Supplemental Figure S2B**) caused a partial rescue of H3K27me3 levels but complete reversal of H3K27ac, possibly due to additive effects of all H3K27 methylation states **(Supplemental Figure S2D)**. These data suggest that polycomb mediated methylation of H3K27 acts as a guardian to prevent hyperacetylation of KSHV episomes.

### 3.3 H3K27 hyperacetylated KSHV episomes show increased basal gene expression and support lytic reactivation by high levels of RTA

Previous work by our group as well as others has shown that KSHV gene expression is modulated by PRC activity (Günther *et al*, 2014; Günther & Grundhoff, 2010; Toth *et al*, 2010) and lytic potential might be increased (Toth *et al*, 2013) in the absence of polycomb function. Indeed, KSHV gene expression of regions, which are decorated with PRC associated histone marks were increased (**Figure 2A)**. This increase in expression, however, was moderate and did not result in spontaneous lytic reactivation. Using RNA-seq we confirmed that the removal of polycomb repression increased PRC target gene expression in the host transcriptome (**Supplemental Figure S2E**). Similarly, broad regions of the KSHV genome showed increased transcriptional activity, especially of genes located in regions, which are characterized by high levels of polycomb repression, including regions of ORF18 to ORF36 and ORF60 to ORF69 **(Figure 2B)**. Surprisingly, this increase in KSHV gene expression did not include a strong increase in lytic markers such as PAN RNA (T1.1) indicating that removing PRC activity did not result in spontaneous lytic reactivation. In line with this observation, we detected increased viral gene expression in polycomb deleted cells using a KSHV BAC16 mutant, which is deficient of RTA (KSHV-BAC16-ΔORF50 (**Figure 2C**), arguing for PRC specific gene regulation independent of lytic reactivation. Similar upregulation of PRC target genes was observed in SLK cells treated with the EZH1/2 inhibitor EPZ-6438 (**Figure 2D**).

**Figure 2:**
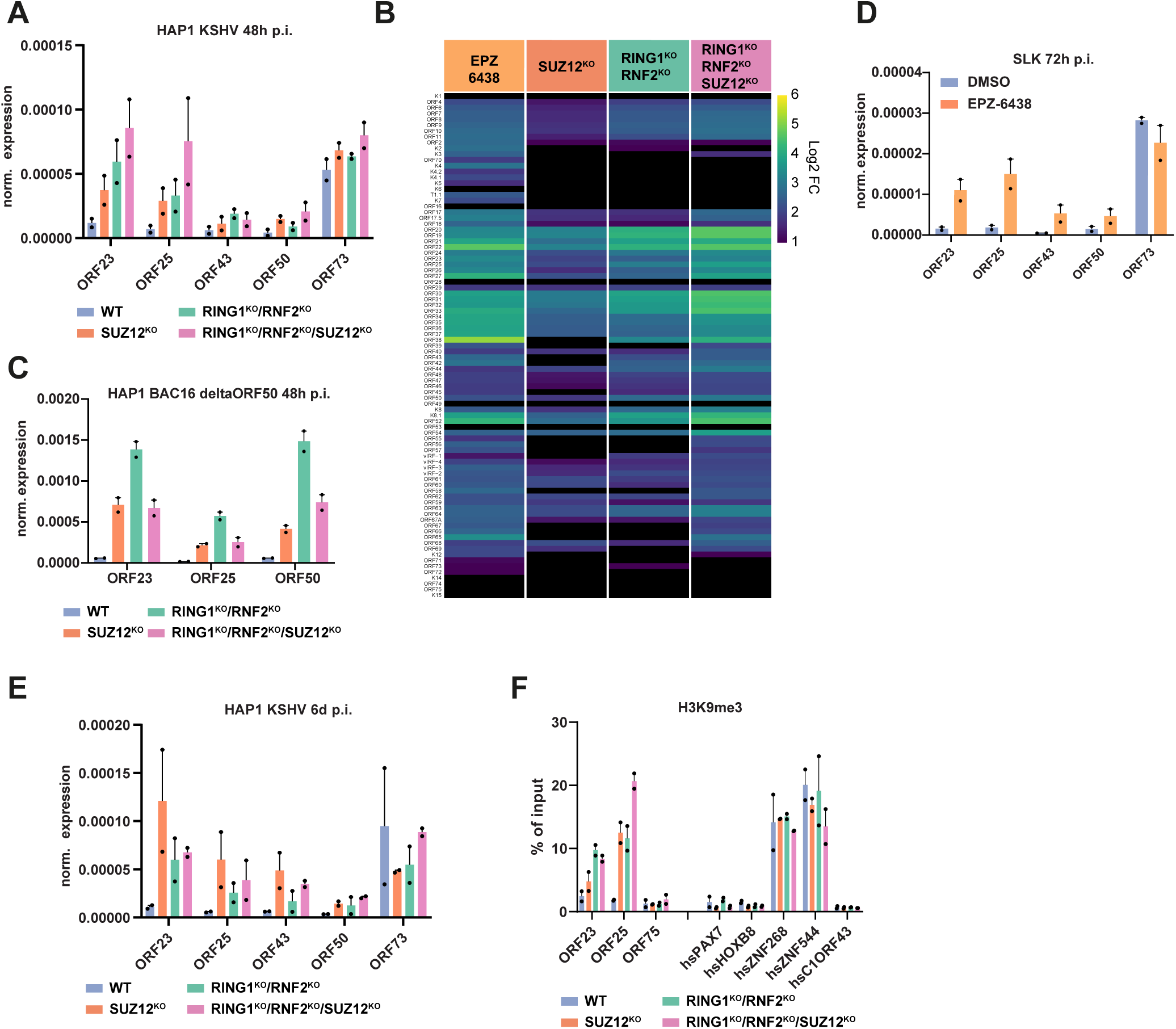
Loss of PRC function causes moderately increased KSHV gene expression, and progressive buildup of H3K9me3 on KSHV episomes. **A)** KSHV gene expression measured by RT-qPCR in HAP1 cells 48h p.i. Expression is normalized to KSHV genomes and human reference genes. (Mean + range; n = two independent biological replicates.) **B)** 48h p.i. RNA-seq of KSHV transcripts in HAP1 DMSO and EPZ-6438 treated as well as different HAP1 PRC knockout cell lines. Significant expression levels changes (P.adj < 0.05) of each KSHV ORF are depicted as color gradient of log2 fold change vs. WT. Nonsignificant changes are depicted as black. **C)** Gene expression of KSHV-BAC16-ΔORF50 in HAP1 cells measured by RT-qPCR cells 48h p.i. Expression is normalized to KSHV genomes and human reference genes. Measuring of ORF50 transcripts was performed by primers targeting the intact ORF50 5’ UTR. (Mean + range; n = two independent biological replicates.) **D)** Treatment of SLK cells with EZH2 inhibitor EPZ-6438 (10 µM) measured by RT-qPCR cells 72h p.i. Expression is normalized to KSHV genomes and human reference genes. (Mean + range; n = two independent biological replicates.) **E)** KSHV gene expression measured by RT-qPCR in HAP1 cells 6 days p.i. Expression is normalized to KSHV genomes and human reference genes. (Mean + range; n = two independent biological replicates.) **F)** ChIP-qPCR analysis of KSHV infected HAP1 cell lines (6d p.i.) H3K9me3. Human control loci (hs) serve as a reference. (Mean + range; n= ChIP replicates of two independent clones

We considered that in the cellular systems used so far, spontaneous lytic reactivation might be limited by insufficient RTA levels and modulation of lytic reactivation by PRC complexes can only be observed in a cellular context that contains spontaneous lytic cell populations. Therefore, we treated BCBL1, as well as *de novo* infected SLK and telomerase-immortalized human microvascular endothelium (TIME) cells with EPZ-6438 for up to 11 days to ensure removal of H3K27me3 from KSHV. Lytic reactivation was not increased in SLK cells. However, after EPZ-6438 treatment of BCBL1 as well as KSHV infected TIME cells we observed an increased fraction of spontaneous lytic cells by expression of ORF59 (**Supplemental Figure S3A**) as well as increased viral genome replication, which could be reversed by addition of replication inhibitor Foscarnet (**Supplemental Figure S3B, S3C**). To complement these observations in HAP1 cells we aimed to overcome insufficient RTA levels by using a doxycycline inducible expression of ORF50/RTA cDNA. Doxycycline inducible RTA expression readily triggers lytic reactivation and is frequently used for virus production of KSHV-BAC constructs (Brulois *et al*, 2012). Indeed, HAP1 cells infected with KSHV and induced with RTA expression show lytic gene expression as well as expression of ORF59 protein (**Supplemental Figure S3C and S3D**). Lytic gene expression, especially of viral genes that are repressed *via* polycomb were increased once PRC function was inhibited. ORF59 positive cell fraction was also moderately increased by PRC deletion (**Supplemental Figure S3D**).

In summary, we have shown that removal of polycomb function leads to a moderately increased basal expression of KSHV genes but did not trigger lytic reactivation. However, PRC deletion modulated lytic gene expression of spontaneous lytic reactivation as well as in cells where lytic cycle entry was induced by RTA overexpression, suggesting that the main function of PRC lytic gene repression is dependent on RTA induction by additional stimuli.

### 3.4 Loss of PRC function causes progressive buildup of H3K9me3 on KSHV episomes

Using our PRC knockout cell lines we could observe that removal of PRC function increases H3K27ac but only moderately increased KSHV gene expression after 48h p.i. (**Figure 2A**) and was not substantially further increased at 6 days post infection (**Figure 2E**) We therefore speculated that viral episomes obtain an altered chromatin composition during continuous propagation of KSHV in PRC deficient cells. Acquisition of alternative heterochromatic histone marks, including H3K9me3, could counteract effects of increased H3K27ac and could explain the lack of increased viral gene expression. Indeed, ChIP-qPCR assays showed a drastic increase of H3K9me3 on KSHV at 6 days p.i., especially once all polycomb function was removed (*RING1^KO^*/*RNF2^KO^*/*SUZ12^KO^*). This was particularly true for regions of strong polycomb recruitment, including ORF23 and ORF25 (**Figure 2F**).

These observations reveal an additional role of polycomb function on KSHV, where the establishment of facultative heterochromatin shields KSHV genomes from constitutive heterochromatin mark H3K9me3.

### 3.5 Increased H3K9me3 de novo deposition is facilitated via the HUSH complex

Recent work investigating the deposition of H3K9me3 on retroviral elements have suggested that the expression of intron-less RNA serves as a marker for recognition of the human silencing hub (HUSH) complex and transcriptional silencing via the methyltransferase SETDB1 (Seczynska *et al*, 2022; Seczynska & Lehner, 2023; Bloor *et al*, 2024). De-repression of intron-less KSHV RNA, caused by removal of PRC function and high H3K27ac levels, could therefore act as a recruitment mechanism of the HUSH transcriptional repressor complex, leading to *de novo* acquisition of H3K9me3. The HUSH transcriptional silencing complex consists of three core subunits, the H3K9me3 binding protein M-phase phosphoprotein 8 (MPP8), the RNA binding Protein periphilin 1 (PPHLN1), and the transcription activation suppressor (TASOR), representing the largest subunit (Müller & Helin, 2024).

To interrogate if HUSH plays a role in H3K9me3 deposition on KSHV we established *TASOR^KO^* HAP1 cells using CRISPR/Cas9. Since strong H3K9me3 deposition is observed in PRC deficient cells we also deleted TASOR in HAP1 cells deficient for PRC2 (*SUZ12^KO^*/*TASOR^KO^*), deficient for PRC1 (*RING1^KO^*/*RNF2^KO^*/*TASOR^KO^*) and for both PRC complexes (*RING1^KO^*/*RNF2^KO^*/*SUZ12^KO^*/*TASOR^KO^*) (**Figure 3A**). Using ChIP-qPCR as well as ChIP-seq we observed a drastic reduction of H3K9me3 once TASOR was deleted, arguing that the HUSH complex is necessary for *de novo* deposition of H3K9me3 after PRC removal (**Figure 3B, 3C and 3D**). Next, we investigated, if HUSH can silence KSHV gene expression via H3K9me3 levels. Therefore, we measured KSHV gene expression via qPCR and RNAseq in PRC and/or TASOR deleted HAP1 cell lines after 7 days post infection. Indeed, removal of PRC complexes together with removal of TASOR strongly increased KSHV expression (**Figure 3E, 3F**). Interestingly, genes that were strongest upregulated once TASOR was removed, are found in regions surrounding ORF18 to ORF36, the exact same location that showed strong enrichment of H3K9me3 in PRC deleted HAP1 cells (**Figure 3D, Supplemental Figure S3E)**. This could indicate, that HUSH binding is initiated by the expression of a specific KSHV transcript within this region.

**Figure 3:**
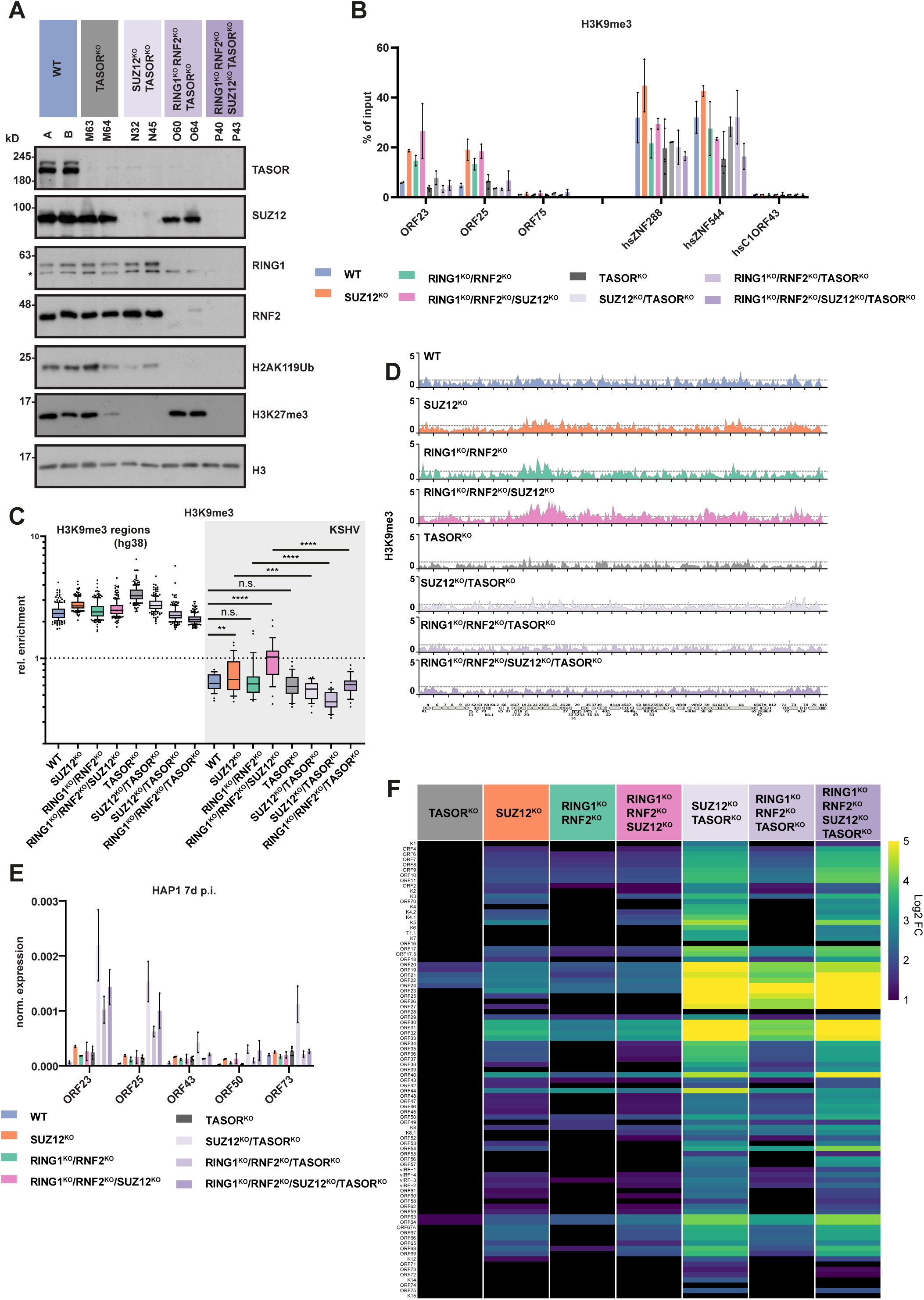
Increased do novo deposition of H3K9me3 is facilitated through the human silencing hub (HUSH) **A)** Immunoblotting of individual HAP1 PRC1 and PRC2 knockout clones in combination with deletion of the HUSH component TASOR. **B)** ChIP-qPCR analysis of KSHV infected HAP1 PRC and HUSH deleted cell lines (7d p.i.) (Mean + range; n= ChIP replicates of two independent clones.) **C**) Quantification of HAP1 KSHV (7d p.i.) H3K9me3 ChIP seq signal in a 10 kb sliding window over the KSHV genome, normalized to negative host regions, sequencing depth and input. Significance of differences of H3K9me3 of viral sequences segments (10kb) were calculated using repeated measure (RM) ANOVA Fisher’s LSD. P > 0.05 (n.s), P ≤ 0.05 (*), P ≤ 0.01 (**), P ≤ 0.001 (***), P ≤ 0.0001 (****) **D)** Read density coverage tracks of H3K9me3 on KSHV (7d p.i.), determined by ChIP-seq (normalized by sequencing depth and input). Shown as fold enrichment over input. Dashed line represents median fold enrichment over input of the host background distribution. **E)** KSHV gene expression measured by RT-qPCR in HAP1 cells 7d p.i. Expression is normalized to KSHV genomes and human reference genes. (Mean + range; n = two independent biological replicates.) **F)** 7d p.i. RNA-seq of KSHV transcripts in HAP1 cells WT as well as different PRC and HUSH knockout cell lines. Significant expression levels changes (P.adj < 0.05) of each KSHV ORF are depicted as color gradient of log2 fold change vs. WT. Nonsignificant changes are depicted as black.

In summary, we observe that removal of PRC function leads to an increase in H3K27ac and increased KSHV gene expression, which can serve as a target for HUSH recruitment and silencing *via* the deposition of H3K9me3.

### 3.6 Loss of Polycomb function leads to failure of episomal maintenance after de novo infection

The fact that loss of PRC function causes an altered chromatin state of KSHV episome, characterized by both increased H3K27ac and increased H3K9me3, represents an intriguing state of metastable latency. We were therefore interested, if other aspects of latent episomes are perturbed. One of the most distinct features of KSHV latency is its persistence in dividing cells through its ability to copy its genome through licensed replication and a LANA mediated tethering mechanism to host chromosomes (Uppal *et al*, 2014). Therefore, we asked if alteration of KSHV chromatin composition could lead to insufficient episomal maintenance after *de novo* infection of KSHV. We monitored episomal content after infection by quantitative PCR in HAP1 WT and PRC knockout cells. Interestingly, we observed an increased episomal loss in *SUZ12^KO^* and an even stronger episomal loss in *RING1^KO^*/*RNF2^KO^* and *RING1^KO^*/*RNF2^KO^*/*SUZ12^KO^* cell lines, indicating that polycomb activity, and especially PRC1 function is a necessary for efficient episomal maintenance (**Figure 4A and Supplemental Figure S4A**). Assuming that episomal loss is progressive over multiple cell divisions, we estimated episomal loss per time as the slope (k) of an exponential growth equation Y=Y0*exp(k*X) fitted to normalized episomes. Episome loss rates increased 4 to 5-fold in PRC deleted cell lines compared to WT HAP1 cells (**Figure 4A**).

**Figure 4:**
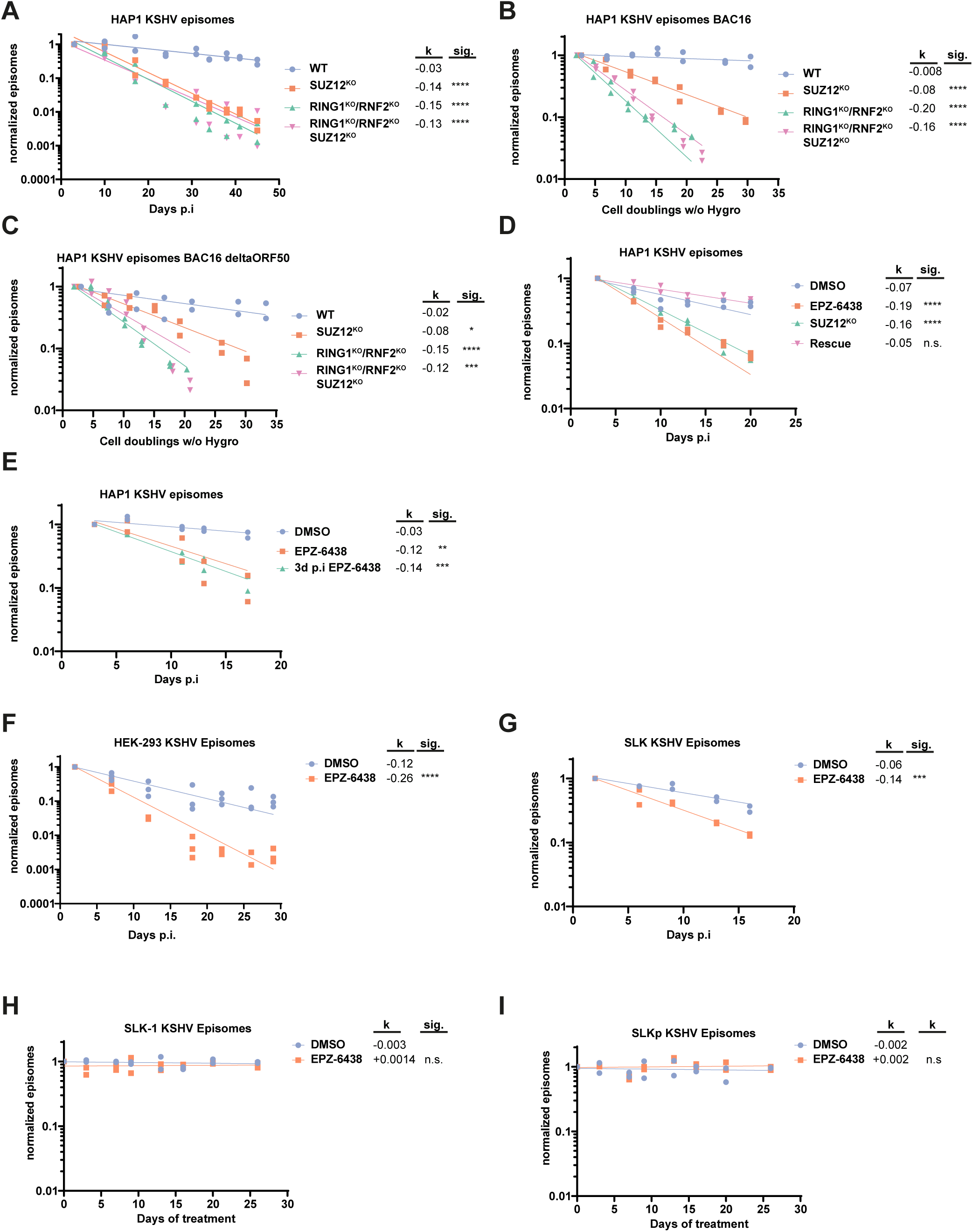
Polycomb function is required for efficient latency establishment. **A)** KSHV episome numbers quantified by quantitative real-time PCR (qPCR) after KSHV de novo infection of HAP1 cell lines, shown as episomes normalized to day 2 p.i. **B)** KSHV-BAC16 and KSHV-BAC16-ΔORF50 **C)** infected cells were selected to achieve 100 % KSHV+ cultures. After removal of hygromycine B following episome quantifications were normalized to Day 0. **D)** KSHV episome numbers quantified by qPCR after KSHV de novo infection of HAP1 cells. HAP1 WT cells were treated with DMSO or 10 µM EPZ-6438 continuously during and after de novo infection. SUZ12^KO^ or Rescue HAP1 cells (SUZ12^KO^ + SUZ12) were infected in parallel with KSHV to compare the dynamic range of EZH2 inhibition affecting episomal maintenance. **E)** KSHV episome numbers quantified by qPCR after KSHV de novo infection of HAP1 cells. HAP1 WT cells were treated with DMSO or 10 µM EPZ-6438 continuously during and after de novo infection (EPZ-6438) or inhibitor treatment was initiated after 72h post infection (3d p.i. EPZ-6438) **F)** KSHV episome numbers quantified by qPCR after KSHV de novo infection of SLK cells. SLK cells were treated with DMSO or 10 µM EPZ-6438 continuously during and after *de novo* infection. **G)** KSHV episome numbers quantified by qPCR after KSHV de novo infection of HEK-293 cells. HEK-293 cells were treated with DMSO or 10 µM EPZ-6438 continuously during and after *de novo* infection. **H and I)** KSHV episome numbers quantified by qPCR in KSHV long-term infected SLK-1 cells (H) and SLKp cells (I). SLK-1 and SLKp cells were continuously treated with DMSO or 10 µM EPZ-6438. Slope of exponential regression (k) is shown next to each sample. Significance (sig.) of difference in slope between control (WT or DMSO) versus treatment (knockout or EPZ-6438) is shown next to each slope (k). P > 0.05 (n.s), P ≤ 0.05 (*), P ≤ 0.01 (**), P ≤ 0.001 (***), P ≤ 0.0001 (****). Cell doublings in 3B and C were calculated according to growth rates depicted in **Supplemental Figure S3F**

Although KSHV gene expression data indicated that this difference in episomal numbers is unlikely caused by alterations in lytic reactivation, we wanted to exclude lytic reactivation by using an RTA deficient KSHV mutant (KSHV-BAC16-ΔORF50). Again, we observed a strong failure in episomal maintenance in PRC knockout cells for both WT and RTA deleted KSHV-BAC16 (**Figure 4B, 4C**). Next, we consolidated this phenotype using pharmacological inhibition of EZH1/2 (EPZ-6438) and rescue experiments. Similar rates of episomal loss for EPZ-6438 treated cells compared to *SUZ12^KO^* were observed (**Figure 4D**) in a dose dependent manner (**Supplemental Figure S4B**), and this was independent if EZH2 inhibition was initiated during infection, or after 3 days, arguing that differences in initial infection rates or altered chromatinization immediately after viral entry are unlikely the cause of inefficient maintenance (**Figure 4E**). The phenotype of increase episomal loss was also recapitulated using quantification of LANA immunofluorescence (**Supplemental Figure S5A, S5B**). Pharmacological inhibition also broadened the range of cell types to investigate this phenotype. Indeed, HEK293 cells (**Figure 4F**) and SLK cells (**Figure 4G**) showed similar differences of episomal maintenance in dependence of EZH2 inhibition, as observed in HAP1 cells. Next, we wanted to test if episomes that have established latency for a prolonged period of time or episomes that have been selected for stable maintenance are sensitive to PRC inhibition. Therefore, we used rKSHV.219 and selected HAP1 WT with puromycin for one month before removing antibiotic selection and treating with EPZ-6438. Again, we could observe that episomal loss was increased by EZH2 inhibition by EPZ-6438 treatment (**Supplemental Figure S4C**). In an effort to extend the observed phenotype of episomal loss to endothelial cells but separate it from the increased spontaneous lytic reactivation after EZH2 inhibitor treatment, we used KSHV-BAC16-ΔORF50 in TIME as well as the endothelial fusion cell line EA.hy926. Both cell lines were selected with hygromycin for one month before removing the selection and treating with EPZ-6438. Again, EZH2 inhibition resulted in increased episomal loss, with a 50 % difference of calculated episomal loss rates (k). (**Supplemental Figure S4D, S4E**). An argument can be made, however, that latency is not fully established within one month after *de novo* infection. In fact important modifications to the KSHV episomes, such as DNA methylation, are established much later (Fröhlich & Grundhoff, 2020) and puromycin selected rKSHV positive cells still lose KSHV episomes spontaneously once the antibiotic selection is released (**Supplemental Figure S4C)**. Therefore, we used SLK-1 and SLKp cells, which are long term infected SLK cell lines that were clonally selected for stable episomal maintenance and therefore show no progressive episomal loss in actively dividing cultures (Grundhoff & Ganem, 2004). Furthermore, they support a strictly latent life cycle with no evidence of spontaneous lytic reactivation. Interestingly, we did not observe any difference in episomal maintenance when SLK-1 or SLKp cells were treated with EPZ-6438 indicating that polycomb function might be dispensable for maintenance of fully matured KSHV episomes (**Figure 4H, 4I**).

Motivated by our observation that H3K9me3 levels were increased in a HUSH dependent manner by loss of PRC function, we asked if episomal loss and function of HUSH were interconnected. Therefore, we infected HAP1 cells devoid of HUSH (*TASOR^KO^*) or lacking both PRC and HUSH activity *SUZ12^KO^*/*TASOR^KO^*, *RING1^KO^*/*RNF2^KO^*/*TASOR^KO^* and *RING1^KO^*/*RNF2^KO^*/*SUZ12^KO^*/*TASOR^KO^*and monitored episomal maintenance. Interestingly, depletion of HUSH had no impact on episomal retention. Similarly, depletion of HUSH in combination with PRC did not rescue the episomal loss phenotype observed by PRC loss alone (**Supplemental Figure S4F**), suggesting that HUSH-mediated heterochromatinization exacerbates but does not initiate episomal loss.

Collectively, we show that polycomb mediated chromatin modifications are important for establishment of episomes that are efficiently maintained after *de novo* infection but fully matured chromatin of KSHV episomes might support stable maintenance independent of polycomb function. Furthermore, increased episomal loss is not a consequence of increased HUSH activity but an attribute directly connected to PRC function.

### 3.7 PRC dependent episomal maintenance is not the result of altered cellular proliferation, latent replication or gross LANA malfunction

The amount of latent viral episomes can be influenced by several physiological processes, which are both regulated by viral as well as cellular molecular pathways. Given that, deletion of polycomb function can influence these processes directly and indirectly we devised a series of experiments to investigate possible contributing mechanisms.

Loss of PRC function combined with KSHV infection could have synergistic effects on cellular proliferation and the observed loss of KSHV positive cells could be caused by a selective growth advantage of KSHV negative cells in a heterogenous population of cells. PRC1 or PRC2 deleted cells showed unaltered growth rates after KSHV infection (**Supplemental Figure S4G**) and episomal loss rates were unchanged once normalized to individual clonal cell proliferation (**Supplemental Figure S4A**). However, Hygromycin selection did alter the growth rates of PRC deleted cell lines after KSHV-BAC16 infection (**Supplemental Figure S4G**). To investigate the effect of cellular proliferation on increased episomal loss, we infected WT and *SUZ12*^KO^ HAP1 cells with BAC16-KSHV and selected with Hygromycin until 100 % KSHV positive cultures were established. Next, we mixed these cultures with KSHV negative cells of the same genotype, marked with BFP, and followed spike-in control cells (BFP), KSHV-positive (GFP) and cells that lost KSHV (colorless) *via* FACS. We observed an increased expansion of KSHV-negative (colorless) cell populations in *SUZ12*^KO^ as well as EPZ-6438 treated cultures. Importantly, the percentage of spike-in control cells within the population did not expand, indicating that KSHV episomes are lost in dividing cells and that increased episomal loss, caused by PRC2 deletion or inhibition, is not resulting from competitive growth advantages (**Figure 5A**). This was in line with the observation that deletion of polycomb complexes did not lead to increased apoptosis in HAP1 cells even when combined with KSHV infection (**Supplemental Figure S5C**). The latency master regulator LANA plays a dual role within episomal maintenance, LANA influences recruitment of the DNA replication machinery during S-Phase and is critical for episomal tethering to mitotic chromosomes (Uppal *et al*, 2014; Barbera *et al*, 2006). We measured LANA gene expression by RT-qPCR and could not detect significant differences between genotypes after 48 h p.i (**Supplemental Figure S5D**). LANA binding to the terminal repeats (TR) measured by ChIP-qPCR assays was also not altered in PRC knockout cell lines after 48 h p.i (**Figure 5B**). Next, we asked if LANA functions are perturbed by polycomb inhibition in a plasmid retention assay (Grundhoff & Ganem, 2003). We transfected plasmids containing eight copies of KSHV terminal repeats and a GFP marker (pGTR8) into LANA overexpressing SLK cells and treated with EPZ-6438 and DMSO as control. By measuring fractions of GFP positive cells over time we could observe that pGTR8 plasmids depleted rapidly in LANA negative cells, whereas LANA could maintain pGTR8 plasmids independent of EZH2 inhibition (**Figure 5C**). Similarly, pGTR8 plasmids were retained with similar efficiency in LANA expressing HAP1 PRC2 knockout cells (**Figure 5D**). Interestingly, calculating episomal loss rates using this assay, PRC1 deletion increased pGTR8 plasmid loss 30 %, indicating that LANA misfunction could contribute to increased KSHV loss in PRC1 depleted cell lines, but cannot explain the drastic increase of KSHV loss (**Figure 4A**).

**Figure 5:**
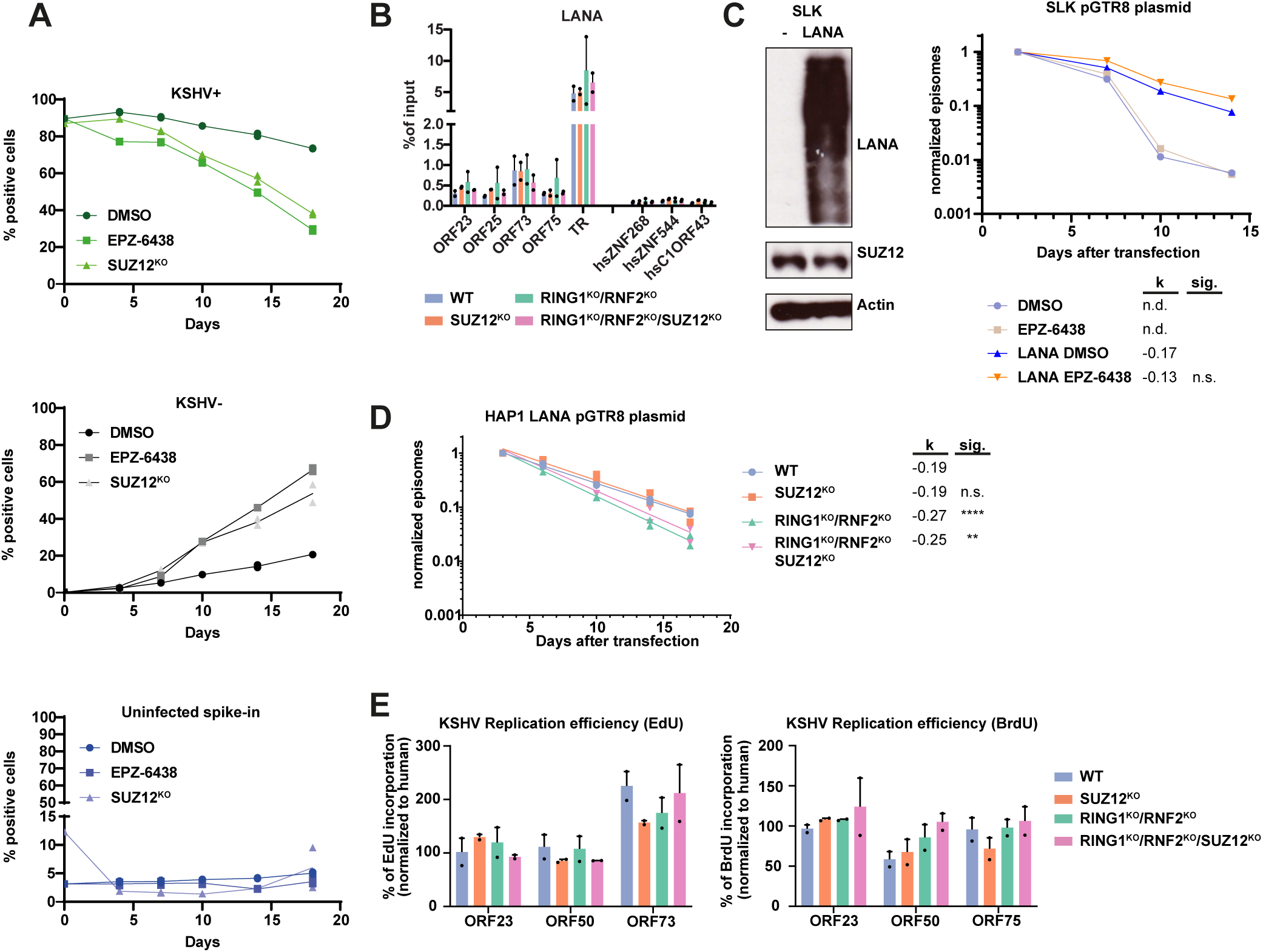
PRC dependent episomal maintenance is not the result of altered cellular proliferation, latent replication or gross LANA malfunction. **A)** Competitive growth assay of cell mixing experiment. KSHV-BAC16 HAP1 cells were mixed with uninfected HAP1 cells in 9:1 ratio. FACS of total population distinguished GFP positive (KSHV+), GFP negative (KSHV-) and BFP positive (uninfected spike-in HAP1 cells) fractions. **B)** LANA binding by ChIP qPCR is detected at KSHV terminal repeats (TR) independent of PRC deletion in HAP1 cells at 48h p.i. **C)** LANA tethered plasmids containing 8 copies of KSHV terminal repeats (pGTR8). Ectopic LANA expression in SLK cells is shown on the left. pGTR8 retention and corresponding slope of exponential regression (k) measured by FACS of GFP positive cell fraction and normalized to first sample (Day 2 after transfection) in DMSO and EPZ-6438 treated SLK and SLK-LANA cells is shown on the right. **D)** pGTR8 retention and corresponding slope of exponential regression (k) measured by FACS of GFP positive cell fraction and normalized to first sample (Day 2 after transfection) in LANA expressing HAP1 cells (n= two independent clones per genotype). Significance (sig.) of difference in slope between control (WT or DMSO) versus treatment (knockout or EPZ-6438) is shown next to each slope (k). P > 0.05 (n.s), P ≤ 0.05 (*), P ≤ 0.01 (**), P ≤ 0.001 (***), P ≤ 0.0001 (****). **E)** Incorporation efficiency of EdU or BrdU after 1h labelling of asynchronously growing HAP1 cells at 48h p.i.

Episomal maintenance is also affected by coordinated replication of the viral genome during S-Phase. Perturbation of latent replication could therefore reduce viral copy numbers. To address the effect of latent replication we measured the amount of nucleotide analog incorporation rates (EdU and BrdU) into viral genomes in asynchronously growing cultures. Here we could not detect differences in incorporation efficiencies relative to human, arguing that latent replication is not affected by loss of polycomb (**Figure 5E**).

Taken together, these data suggest that loss of polycomb function progressively increases episomal loss in dividing cells after de novo infection. This loss is not caused by altered cellular proliferation and incorporation rates of nucleotide analogs indicate no change in latent replication. Furthermore, basic functionality of LANA is unperturbed in PRC deleted cell lines, indicating that increased episomal loss is the consequence of altered KSHV episomes, rather than an altered cellular state, which could favor loss of extrachromosomal DNA content. This is in line with the fact that episomal loss is not induced in SLKp and SLK-1 cells with established KSHV latency (**Figure 4H, 4I**).

## 4. Discussion

The decoration of viral genomes with histones and its corresponding modifications after *de novo* infections represents a crucial stage of latency establishment and a transition of “naked” viral DNA towards a genomic unit with well-defined transcriptional output. In this study, we have systematically deleted core members of the polycomb complexes including simultaneous deletion of both PRC1 and PRC2. While we find that PRC1 and PRC2 can act on KSHV independently, the fact that H3K27me3 levels were strongly reduced by loss of PRC1 argues that the dominant axis of PRC recruitment on KSHV occurs *via* variant PRC1 complexes.

Polycomb repressive complexes have been reported as a block of KSHV lytic reactivation (Günther & Grundhoff, 2010; Toth *et al*, 2010, 2013; Günther *et al*, 2014; Naik *et al*, 2020). Indeed, we detect an impact of PRC function on lytic reactivation. However, this was only true in conditions where KSHV ORF50 (RTA) was elevated, either using inhibitors in KSHV infected cell lines that show high rates of spontaneous lytic reactivation or by forced expression of RTA in cell lines in which KSHV maintains a strictly latent lifecycle. Although PRC complexes can act directly on the KSHV ORF50 promoter (Günther & Grundhoff, 2010; Toth *et al*, 2013) we argue that the main function of PRC function on repression of lytic genes is downstream of RTA.

We and others have observed that H3K27me3 is the dominant repressive mark, detectable 24h p.i and covering large portions of the KSHV genome 72h p.i., at levels that are on par with human positive control regions (Günther & Grundhoff, 2010; Toth *et al*, 2010). H3K9me3 deposition, however, can only be detected at low levels on KSHV, compared to host positive controls during *de novo* infection in wildtype cells and is only enrich on KSHV in cell cultures that have been infected with KSHV for an extended period of time, such as BCBL1 or SLKp cells. Furthermore, this H3K9me3 acquisition is focused around two specific regions of late genes (ORF19-ORF25) and (ORF64-ORF67) (Günther & Grundhoff, 2010). Interestingly, these regions are exactly the same, which we see decorated by H3K9me3 once PRC functions are impaired, early after de novo infection. This argues that an intrinsic feature within these regions can act as a signal for H3K9me3 buildup. Our demonstration, that the H3K9me3 levels are dependent on HUSH function indicate that this signal could be the expression of an intron-less RNA originating within this genomic area. As H3K9me3 and H3K27ac are assumed to decorate mutual exclusive sites in the human genome, it is arguably unlikely that those two marks are located at the same viral episomes. Given that ChIP-seq interrogates the bulk population of KSHV genomes, it is possible that different chromatin configurations of viral episomes co-exist within different cells or even within the same host cell. This could mean that hyperacetylation of H3K27, increased viral gene transcription, and buildup of H3K9me3 are functionally linked and occur in a stepwise progression. The capability of the HUSH complex to bind intron-less genes and our observation that removal of HUSH diminishes H3K9me3 agrees with this model. However, a potential competition between PRC bound chromatin and the proposed RNA independent recruitment of HUSH via chromatin binding protein NP220 could explain HUSH dependent H3K9me3 deposition without the need for a specific RNA feature within these specific regions within the KSHV genome (Zhu *et al*, 2018). Here we propose that PRC complexes, not only ensure gene repression on KSHV but also protect from detection of the HUSH complex and its potential antiviral repressive function (Roy & Ghosh, 2024).

Interestingly, Roubille and colleagues (Roubille *et al*, 2024) recently reported HUSH-mediated H3K9me3 deposition on PML NB-associated genomes of a quiescent-prone HSV-1 mutant. Since KSHV genomes do not associate with PML NB during latent de novo infection (Günther *et al*, 2019), the mechanisms that attract the HUSH complex to KSHV and HSV-1 may differ. While Roubille et al. furthermore observed H3K9me3 deposition across the entire HSV-1 genome, we observed buildup preferentially in sub-genomic regions of KSHV that show substantial dysregulation in the absence of PRCs. The fact that the same regions are marked by H3K9me3 in tumor-derived BCBL1 cells suggest that the same, presumably transcription-dependent HUSH recruitment mechanisms act upon long-term infected cells *in vivo*. In agreement with the protective role of PRC complexes suggested by our study, H3K27me3 and H3K9me3 are furthermore clearly anti-correlated in BCBL1 cells (Günther & Grundhoff, 2010). Finally, H3K9me3 patterns in SLKp cells are similar to those in BCBL1 cells but show considerably lower overall enrichment, suggesting that a smaller fraction of episomes is decorated by H3K9me3 in such cells. Collectively, our data therefore suggest a model in which episodic firing of lytic promoters leads to a slow exchange of facultative against constitutive heterochromatin marks. If and why promoters in the observed H3K9me3-marked regions may be especially prone to sporadic firing remains to be determined.

After *de novo* infection and during latency establishment, KSHV episomes face several challenges in order to be retained in dividing cells. Previous reports, characterizing the efficiency of episome retention, show that KSHV genomes are vulnerable to episomal loss after de novo infection and are inefficiently maintained in dividing cultures (Grundhoff & Ganem, 2004). In fact, only a minority of cells remain KSHV positive after *de novo* infection. Here we report, that PRC mediated chromatin composition might play an important role during latency establishment after de novo infection. We propose that PRC function protects incoming KSHV episomes during a vulnerable phase of latency establishment, providing the means to enforce transcriptional regulation of viral genes, as well as shielding viral genomes from constitutive heterochromatin and episomal loss.

However, the exact mechanism how viral episomes are lost during cell division remains unknown. We have excluded the possibilities of cellular proliferation, latent replication and gross LANA malfunction. Furthermore, the fact that polycomb function is crucial only during latency establishment early after *de novo* infection, but is dispensable for episomal maintenance of established KSHV latent episomes, argues that episomal loss is the consequence of altered KSHV episomes, rather than deregulation of cellular genes or host chromatin composition. It is tempting to speculate that episomal loss occurs during mitosis, as defects during tethering of viral genomes to host chromosomes would result in mis-segregation of KSHV episomes to the host cytoplasm, which would likely lead to the elimination of viral DNA. However, the question remains, how altered KSHV chromatin could influence tethering of viral genomes to host chromosomes. KSHV LANA tethers the long stretch of KSHV terminal repeats to nucleosomes (Barbera *et al*, 2006), this includes nucleosomes bound to host DNA but also additional KSHV genomes. This has led to the proposal that episome clustering is an important feature of KSHV episomal maintenance. The aberrant acquisition of H3K27ac and increased viral transcription on viral episomes could influence compaction and cluster formation, thereby weakening episomal tethering.

Collectively, we have shown that PRC function is a central mechanism in KSHV latency establishment ensuring transcriptional repression of viral genes. We expand the current view of PRC function on KSHV and propose that polycomb can protect the viral genome during a vulnerable phase early after *de novo* infection from acquisition of constitutive heterochromatin and episomal loss.

## Acknowledgements

We thank the members of the labs of A. Grundhoff and N. Fischer for discussion and technical advice. We thank the LIV technology platforms Flow Cytometry/FACS and High-Throughput Sequencing as well as Jaqueline Nakel and Armin Günther for technical support. This work was supported by the Deutsche Forschungsgemeinschaft (DFG, German Research Foundation) in the framework of the Research Unit FOR5200 DEEP-DV (443644894) through project GR 3318/5-1.

## Figure legends

**Supplemental Figure S1:**
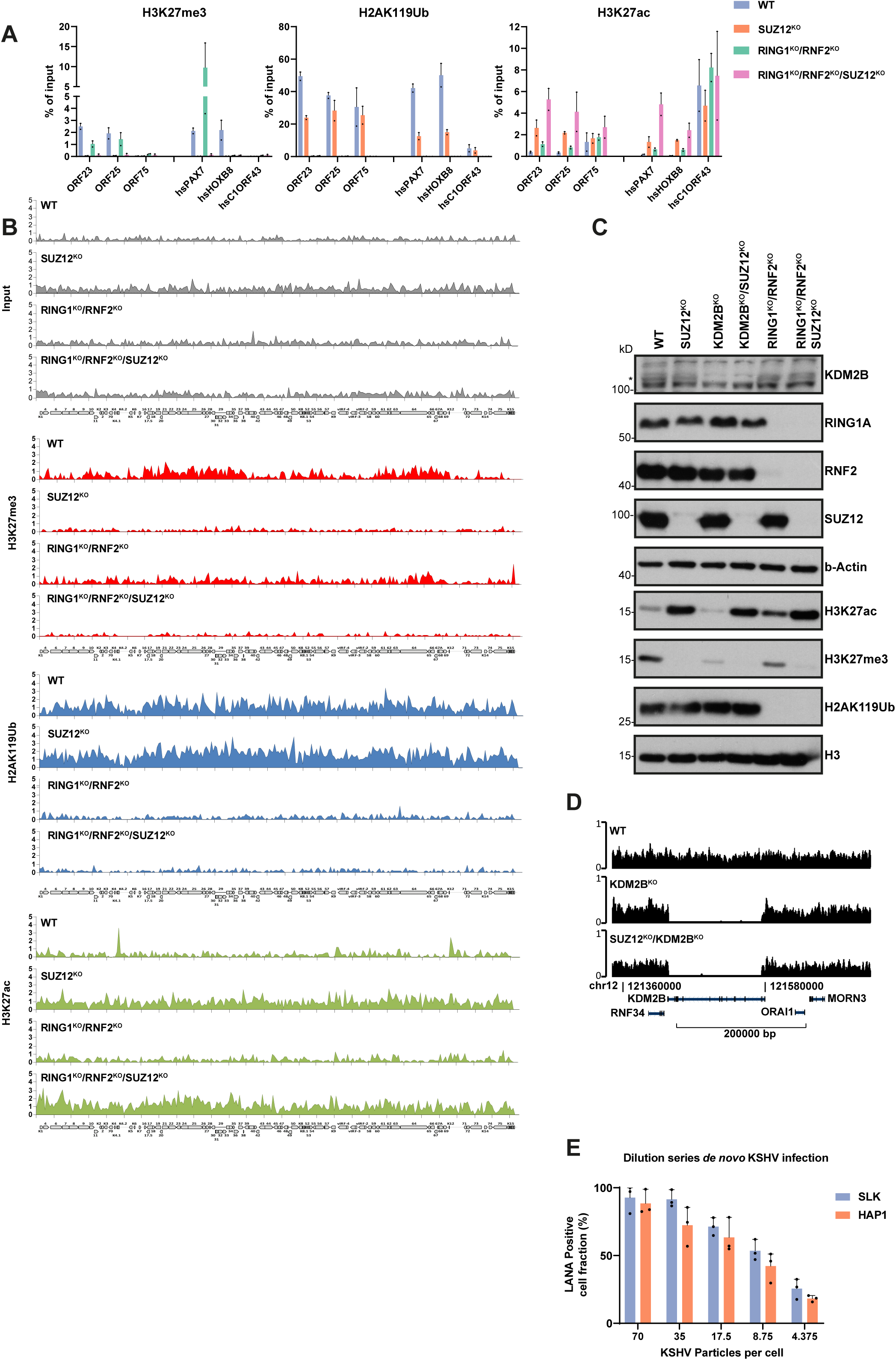
**A)** ChIP-qPCR analysis of KSHV infected HAP1 cell lines (48h p.i.) Investigated mark or factor is shown on top of each graph. Human control loci (hs) serve as a reference. (Mean + range; n= two independent ChIP replicates.) **B)** Read density coverage tracks of histone marks on KSHV, determined by ChIP-seq. **C)** Immunoblot analysis of HAP1 knockout clonal cell lines. Individual or combinatorial deletion of genes is indicated at the top of each blot. KDM2B band deleted in *KDM2B^KO^* cells is indicated with an asterisk symbol (*). **D)** Genome browser view of the locus containing *KDM2B* in *WT*, *KDM2B*^KO^ and *SUZ12*^KO^*/KDM2B*^KO^ HAP1 cells, showing complete removal of the locus. **E)** Quantification of LANA IF (48 h p.i.) of dilution series of *de novo* KSHV infection of 4×10^4^ SLK and HAP1 cells with indicated number of infectious particles (Mean + range; n= 3 independent infections).

**Supplemental Figure S2:**
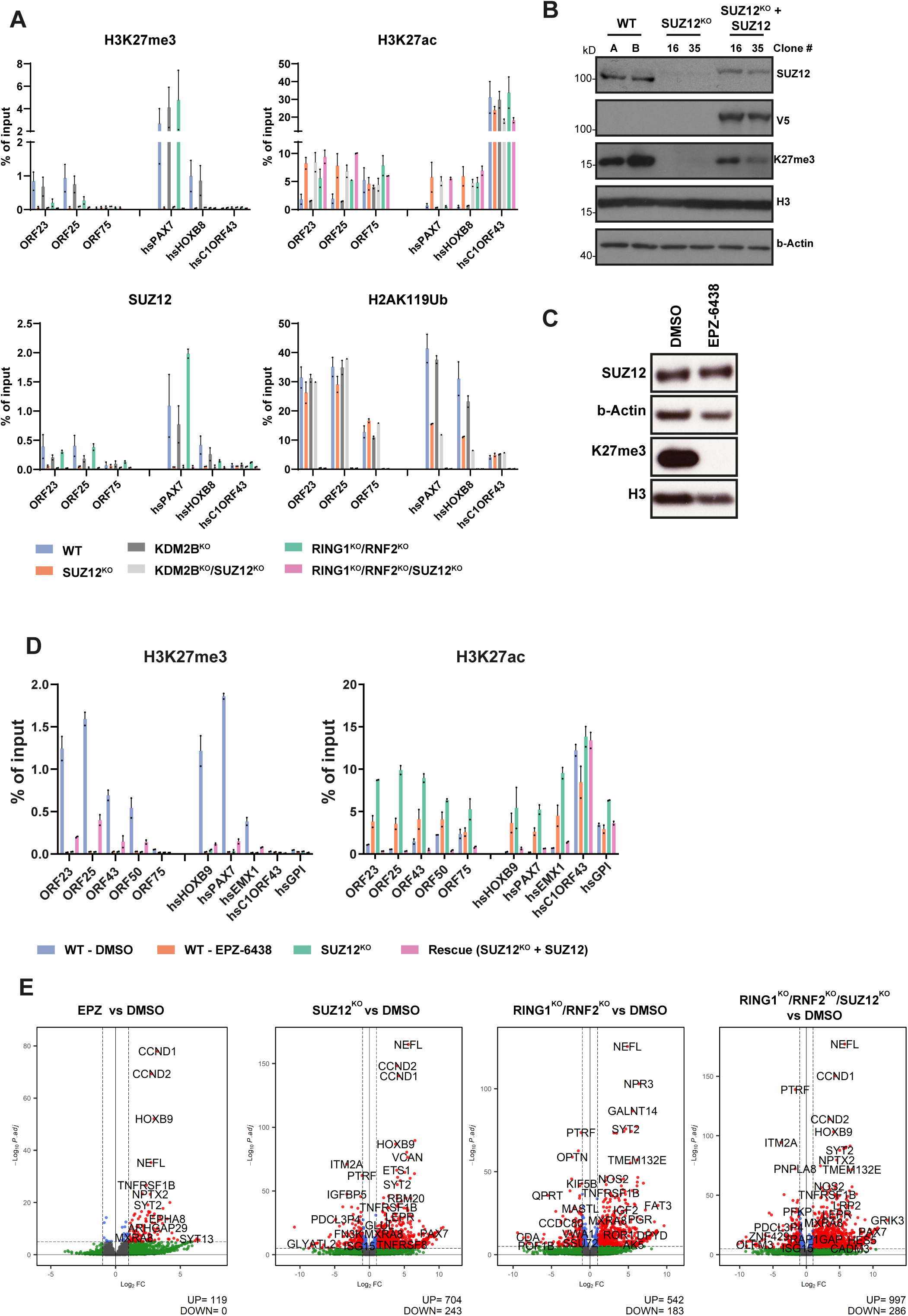
**A)** ChIP-qPCR analysis of KSHV infected HAP1 cell lines (48h p.i.) Investigated mark or factor is shown on top of each graph. Human control loci (hs) serve as a reference. (Mean + range; n= two independent ChIP replicates.) **B)** Immunoblotting of SUZ12 recue cells **C)** EZH2 Inhibitor treatment (5 days, 10 µM EPZ-6438) of HAP1 cells shows loss of H3K27me3. **D)** ChIP-qPCR analysis of KSHV infected HAP1 cells (48h p.i.) after DMSO (WT-DMSO), 10 µM EPZ-6438 treatment (WT-EPZ-6438), SUZ12^KO^, or Rescue HAP1 cells (SUZ12^KO^ + SUZ12). Investigated mark is shown on top of each graph. Human control loci (hs) serve as a reference. (Mean + range; n= two independent ChIP replicates.) **E)** Differential expression of human transcriptome of HAP1 cells DMSO or EPZ6438 treated as well as different PRC knockout cell lines 48h p.i.. Number of significantly changed genes is shown below. (UP: log2FoldChange >= 1, padj < 10e-6; DOWN: log2FoldChange <= -1, padj < 10e-6).

**Supplemental Figure S3:**
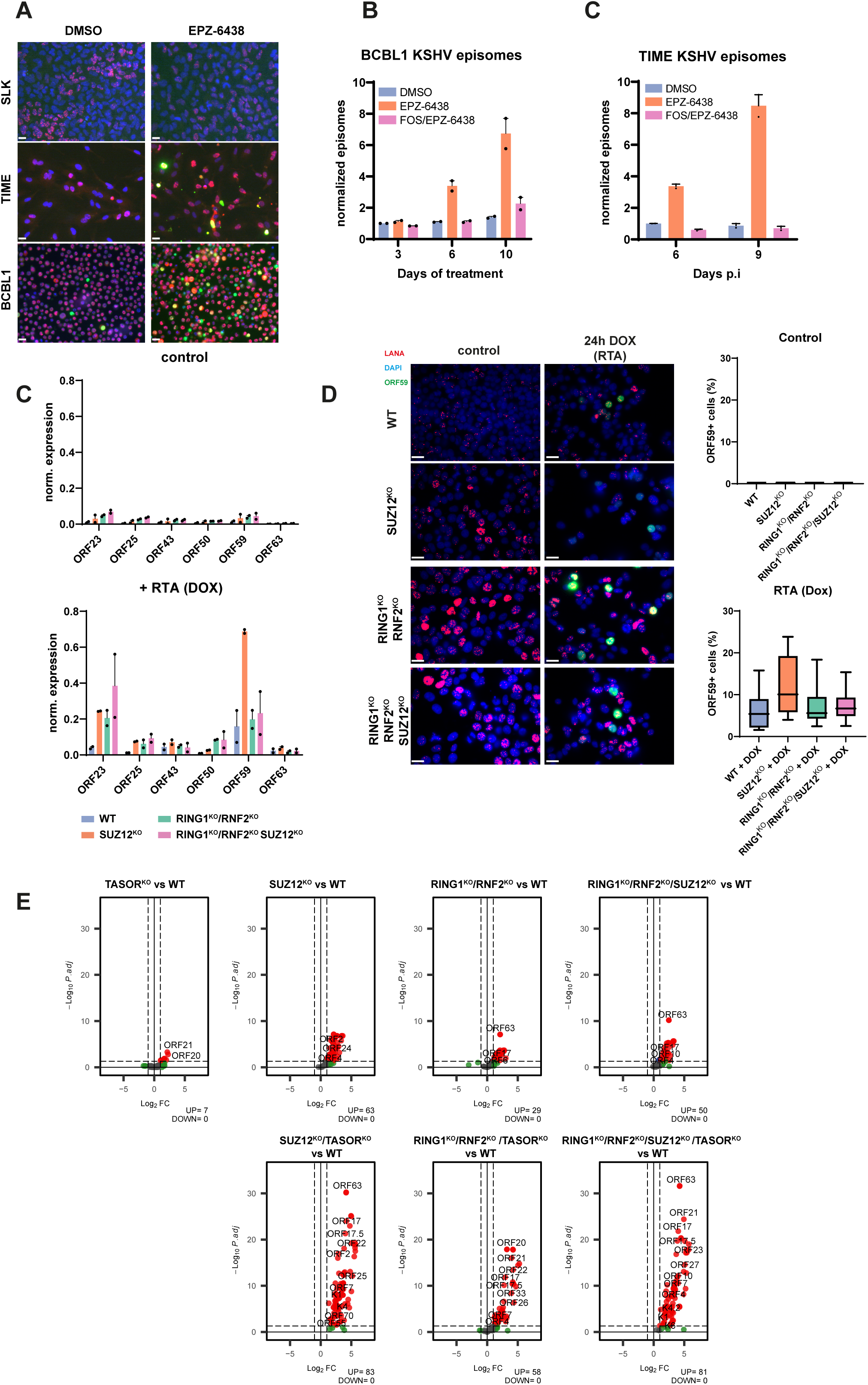
**A)** EPZ-6438 treatment triggers lytic reactivation in BCBL1 and TIME but not SLK cells. Immunofluorescence of LANA and ORF59 in SLK or TIME cells after *de novo* infection and DMSO or 10µM EPZ-6438 treatment (6 days post infection) or 11 days of treatment (BCBL1). Scale bar: 20 µm **B)** KSHV episome numbers quantified by qPCR in BCBL1 after 6 and 9 days of treatment with 10µM EPZ-6438 alone or in combination with 400 µM foscarnet (FOS). **C)** KSHV episome numbers quantified by qPCR in TIME cells infected with KSHV after 6 and 9 days treated with 10 µM EPZ-6438 alone or in combination with 400 µM foscarnet (FOS). **C)** Gene expression measured by RT-qPCR of HAP1 cells 48h post infection treated with control or 1 µg/ml doxycycline (DOX). Expression is normalized to KSHV genomes and human reference genes. (Mean + range; n = two independent biological replicates.) **D)** ORF59 expression measured by IF of HAP1 cells 48h post infection treated with control or 1 µg/ml doxycycline (DOX) Scale bar: 20 µm. Average ORF59+ cell fraction is shown adjacent. Positive cell fractions were pooled from eight images (four images per clone of respective genotype). **E)** Differential expression of KSHV ORFs of HAP1 cells WT and different PRC and HUSH knockout cell lines 6 days p.i.. Number of significantly changed genes is shown below. (UP: log2FoldChange >= 1, padj < 0.05; DOWN: log2FoldChange <= -1, padj < 0.05).

**Supplemental Figure S4.**
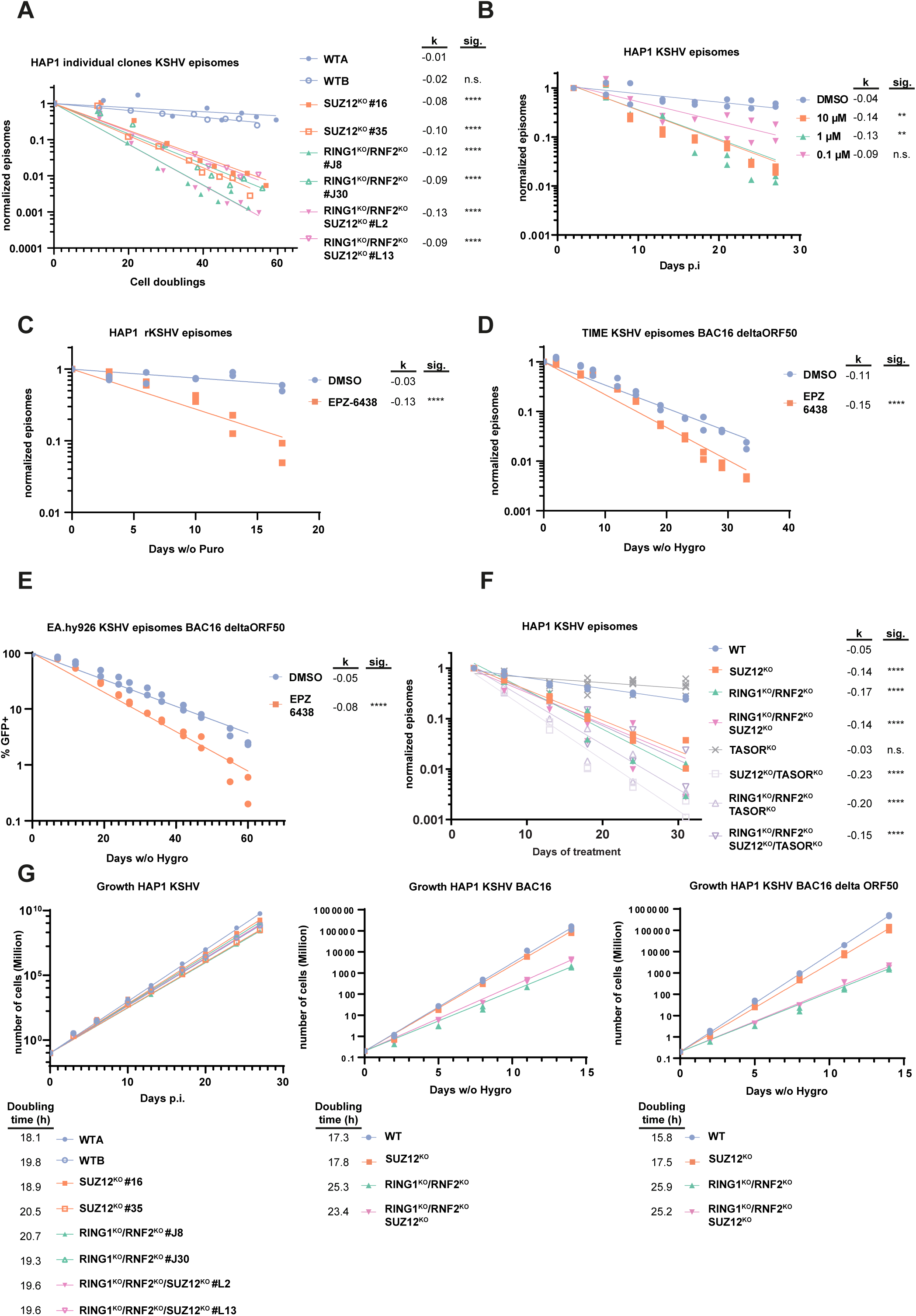
**A)** KSHV episome numbers quantified by quantitative real-time PCR (qPCR) after KSHV de novo infection of HAP1 cell lines, shown as episomes normalized to day 2 p.i. for each individual cellular clone used in **Figure 3A**. **B)** KSHV episome numbers quantified by qPCR after KSHV de novo infection of HAP1 cell lines treated with indicated amounts of EPZ-6438, shown as episomes normalized to day 2 p.i. **C)** KSHV episome numbers quantified by qPCR of HAP1 WT cells infected with rKSHV.219 selected with puromycin and treated with 10µM of EPZ-6438 without addition of puromycin, shown as episomes normalized to day 0 of treatment. **D)** KSHV episome numbers quantified by qPCR of TIME cells infected with KSHV-BAC16-ΔORF50 selected with hygromycin and treated with 10µM of EPZ-6438 without addition of hygromycin, shown as episomes normalized to day 0 of treatment **E)** KSHV positive cells quantified by FACS (GFP+) infected with KSHV-BAC16-ΔORF50 selected with hygromycin and treated with 10µM of EPZ-6438 without addition of hygromycin. **F)** KSHV episome numbers quantified by quantitative real-time PCR (qPCR) after KSHV de novo infection of HAP1 cell lines, shown as episomes normalized to day 2 p.i. Slope of exponential regression (k) is shown next to each sample. Significance (sig.) of difference in slope between control (WT or DMSO) versus treatment (knockout or EPZ-6438) is shown next to each slope (k). P > 0.05 (n.s), P ≤ 0.05 (*), P ≤ 0.01 (**), P ≤ 0.001 (***), P ≤ 0.0001 (****). **G)** HAP1 growth rates after infection with KSHV, KSHV-BAC16 and KSHV-BAC16-ΔORF50. Cell doublings in S4A were calculated according to growth rates depicted in **Supplemental Figure S4G.**

**Supplemental Figure S5:**
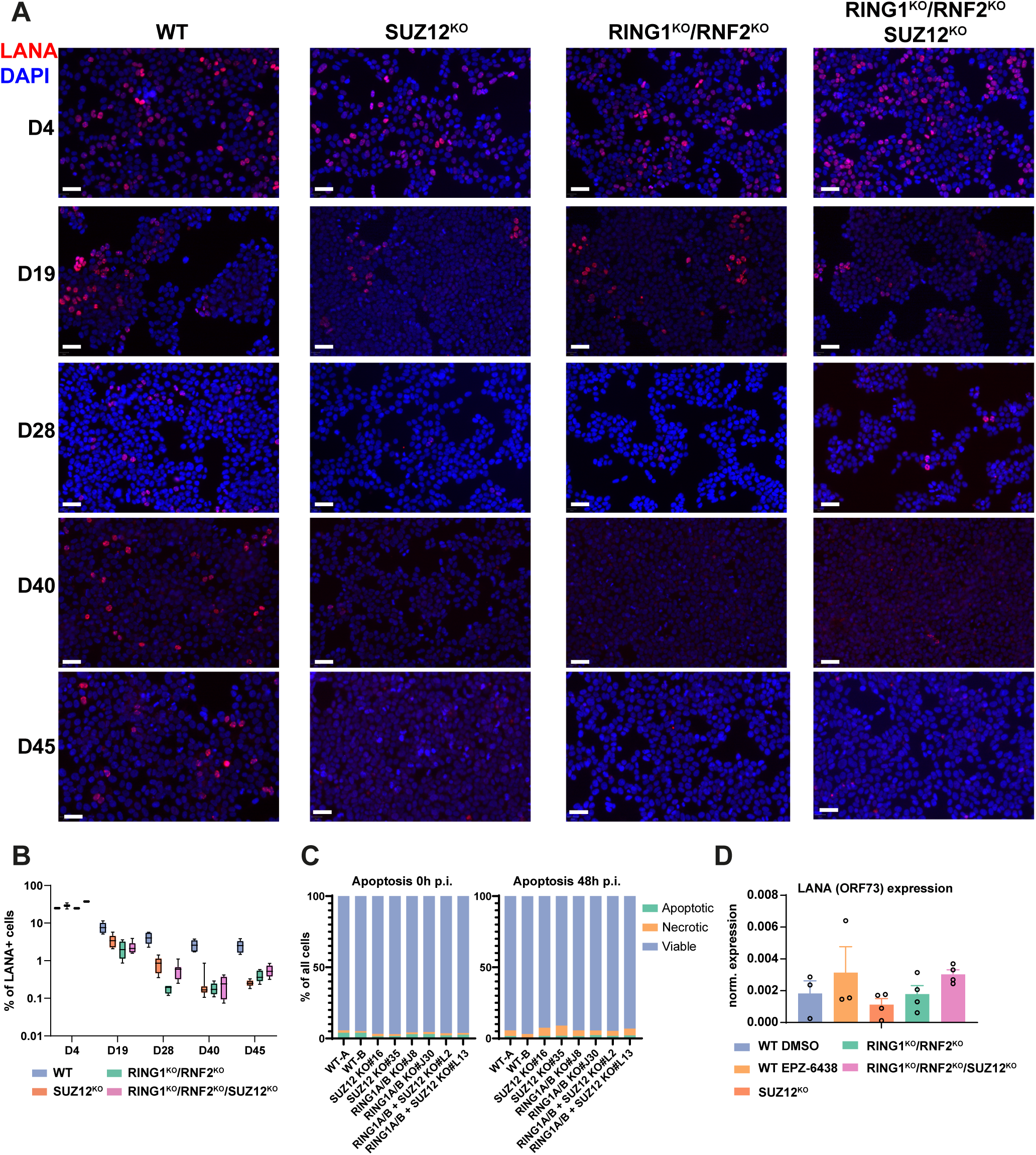
**A)** KSHV+ cell detection by LANA immunostaining of HAP1 cell. Representative images **B)** Automated quantification of LANA immunostaining of HAP1 cells. Two clones per genotype and four pictures per clonal cell line was analyzed. Median counted cells per genotype per time point: n=45000. **C)** Quantification of cellular viability and apoptosis rates of uninfected and KSHV infected (48h p.i.) HAP1 cell lines **D)** RT qPCR of KSHV ORF73 (LANA) at 48h p.i. (n= 4, two biological replicates of two independent clonal HAP1 lines)

## Notes

### Competing Interest Statement

The authors have declared no competing interest.

## References

Bankhead P, Loughrey MB, Fernández JA, Dombrowski Y, McArt DG, Dunne PD, McQuaid S, Gray RT, Murray LJ, Coleman HG, et al (2017) QuPath: Open source software for digital pathology image analysis. Sci Rep 7: 16878

Barbera AJ, Chodaparambil J V., Kelley-Clarke B, Joukov V, Walter JC, Luger K & Kaye KM (2006) The Nucleosomal Surface as a Docking Station for Kaposi’s Sarcoma Herpesvirus LANA. Science (80-) 311: 856–861

Beringer M, Pisano P, Di Carlo V, Blanco E, Chammas P, Vizán P, Gutiérrez A, Aranda S, Payer B, Wierer M, et al (2016) EPOP Functionally Links Elongin and Polycomb in Pluripotent Stem Cells. Mol Cell 64: 645–658

Blackledge NP & Klose RJ (2021) The molecular principles of gene regulation by Polycomb repressive complexes. Nat Rev Mol Cell Biol 22: 815–833

Bloor S, Wit N & Lehner PJ (2024) RNA binding by Periphilin plays an essential role in initiating silencing by the HUSH complex. Nucleic Acids Res 15: 9492

Bracken AP, Dietrich N, Pasini D, Hansen KH & Helin K (2006) Genome-wide mapping of polycomb target genes unravels their roles in cell fate transitions. Genes Dev 20: 1123–1136

Broussard G & Damania B (2020) Regulation of KSHV Latency and Lytic Reactivation. Viruses 12: 1034

Brulois KF, Chang H, Lee AS-Y, Ensser A, Wong L-Y, Toth Z, Lee SH, Lee H-R, Myoung J, Ganem D, et al (2012) Construction and Manipulation of a New Kaposi’s Sarcoma-Associated Herpesvirus Bacterial Artificial Chromosome Clone. J Virol 86: 9708–9720

Cao R, Wang L, Wang H, Xia L, Erdjument-Bromage H, Tempst P, Jones RS & Zhang Y (2002) Role of Histone H3 Lysine 27 Methylation in Polycomb-Group Silencing. Science (80-) 298: 1039–1043

Carette JE, Raaben M, Wong AC, Herbert AS, Obernosterer G, Mulherkar N, Kuehne AI, Kranzusch PJ, Griffin AM, Ruthel G, et al (2011) Ebola virus entry requires the cholesterol transporter Niemann– Pick C1. Nature 477: 340–343

Cesarman E, Chang Y, Moore PS, Said JW & Knowles DM (1995) Kaposi’s Sarcoma–Associated Herpesvirus-Like DNA Sequences in AIDS-Related Body-Cavity–Based Lymphomas. N Engl J Med 332: 1186–1191

Dittmer DP & Damania B (2019) Kaposi’s Sarcoma-Associated Herpesvirus (KSHV)-Associated Disease in the AIDS Patient: An Update. In Cancer Treatment and Research pp 63–80.

Dobin A, Davis CA, Schlesinger F, Drenkow J, Zaleski C, Jha S, Batut P, Chaisson M & Gingeras TR (2013) STAR: Ultrafast universal RNA-seq aligner. Bioinformatics 29: 15–21

Farcas AM, Blackledge NP, Sudbery I, Long HK, McGouran JF, Rose NR, Lee S, Sims D, Cerase A, Sheahan TW, et al (2012) KDM2B links the polycomb repressive complex 1 (PRC1) to recognition of CpG islands. Elife 2012: 1–26

Ferrari KJ, Scelfo A, Jammula S, Cuomo A, Barozzi I, Stützer A, Fischle W, Bonaldi T & Pasini D (2014) Polycomb-Dependent H3K27me1 and H3K27me2 Regulate Active Transcription and Enhancer Fidelity. Mol Cell 53: 49–62

Fröhlich J & Grundhoff A (2020) Epigenetic control in Kaposi sarcoma-associated herpesvirus infection and associated disease. Semin Immunopathol 42: 143–157

Fursova NA, Turberfield AH, Blackledge NP, Findlater EL, Lastuvkova A, Huseyin MK, Dobrinić P & Klose RJ (2021) BAP1 constrains pervasive H2AK119ub1 to control the transcriptional potential of the genome. Genes Dev 35: 749–770

Gao Z, Zhang J, Bonasio R, Strino F, Sawai A, Parisi F, Kluger Y & Reinberg D (2012) PCGF Homologs, CBX Proteins, and RYBP Define Functionally Distinct PRC1 Family Complexes. Mol Cell 45: 344–356

Glancy E, Ciferri C & Bracken AP (2021) Structural basis for PRC2 engagement with chromatin. Curr Opin Struct Biol 67: 135–144

Grundhoff A & Ganem D (2003) The Latency-Associated Nuclear Antigen of Kaposi’s Sarcoma-Associated Herpesvirus Permits Replication of Terminal Repeat-Containing Plasmids. J Virol 77: 4470–4470

Grundhoff A & Ganem D (2004) Inefficient establishment of KSHV latency suggests an additional role for continued lytic replication in Kaposi sarcoma pathogenesis. J Clin Invest 113: 124–136

Günther T, Fröhlich J, Herrde C, Ohno S, Burkhardt L, Adler H & Grundhoff A (2019) A comparative epigenome analysis of gammaherpesviruses suggests cis-acting sequence features as critical mediators of rapid polycomb recruitment. PLOS Pathog 15: e1007838

Günther T & Grundhoff A (2010) The Epigenetic Landscape of Latent Kaposi Sarcoma-Associated Herpesvirus Genomes. PLoS Pathog 6: e1000935

Günther T, Schreiner S, Dobner T, Tessmer U & Grundhoff A (2014) Influence of ND10 Components on Epigenetic Determinants of Early KSHV Latency Establishment. PLoS Pathog 10: e1004274

Højfeldt JW, Laugesen A, Willumsen BM, Damhofer H, Hedehus L, Tvardovskiy A, Mohammad F, Jensen ON & Helin K (2018) Accurate H3K27 methylation can be established de novo by SUZ12-directed PRC2. Nat Struct Mol Biol 25: 225–232

Hopcraft SE, Pattenden SG, James LI, Frye S, Dittmer DP & Damania B (2018) Chromatin remodeling controls Kaposi’s sarcoma-associated herpesvirus reactivation from latency. PLOS Pathog 14: e1007267

Juillard F, Tan M, Li S & Kaye KM (2016) Kaposi’s Sarcoma Herpesvirus Genome Persistence. Front Microbiol 7: 1–15

Kasinath V, Beck C, Sauer P, Poepsel S, Kosmatka J, Faini M, Toso D, Aebersold R & Nogales E (2021) JARID2 and AEBP2 regulate PRC2 in the presence of H2AK119ub1 and other histone modifications. Science (80-) 371

Knutson SK, Warholic NM, Wigle TJ, Klaus CR, Allain CJ, Raimondi A, Porter Scott M, Chesworth R, Moyer MP, Copeland RA, et al (2013) Durable tumor regression in genetically altered malignant rhabdoid tumors by inhibition of methyltransferase EZH2. Proc Natl Acad Sci 110: 7922–7927

Langmead B, Trapnell C, Pop M & Salzberg SL (2009) Ultrafast and memory-efficient alignment of short DNA sequences to the human genome. Genome Biol 10: R25

Laugesen A, Højfeldt JW & Helin K (2019) Molecular Mechanisms Directing PRC2 Recruitment and H3K27 Methylation. Mol Cell 74: 8–18

Li H, Liefke R, Jiang J, Kurland JV, Tian W, Deng P, Zhang W, He Q, Patel DJ, Bulyk ML, et al (2017) Polycomb-like proteins link the PRC2 complex to CpG islands. Nature 549: 287–291

Liao Y, Smyth GK & Shi W (2014) FeatureCounts: An efficient general purpose program for assigning sequence reads to genomic features. Bioinformatics 30: 923–930

Love MI, Huber W & Anders S (2014) Moderated estimation of fold change and dispersion for RNA-seq data with DESeq2. Genome Biol 15: 550

Margueron R, Li G, Sarma K, Blais A, Zavadil J, Woodcock CL, Dynlacht BD & Reinberg D (2008) Ezh1 and Ezh2 Maintain Repressive Chromatin through Different Mechanisms. Mol Cell 32: 503–518

Müller I & Helin K (2024) Keep quiet: the HUSH complex in transcriptional silencing and disease. Nat Struct Mol Biol 31: 11–22

Naik NG, Nguyen TH, Roberts L, Fischer LT, Glickman K, Golas G, Papp B & Toth Z (2020) Epigenetic factor siRNA screen during primary KSHV infection identifies novel host restriction factors for the lytic cycle of KSHV. PLOS Pathog 16: e1008268

Piunti A & Shilatifard A (2021) The roles of Polycomb repressive complexes in mammalian development and cancer. Nat Rev Mol Cell Biol 22: 326–345

Poepsel S, Kasinath V & Nogales E (2018) Cryo-EM structures of PRC2 simultaneously engaged with two functionally distinct nucleosomes. Nat Struct Mol Biol 25: 154–162

Riising EM, Comet I, Leblanc B, Wu X, Johansen JV & Helin K (2014) Gene silencing triggers polycomb repressive complex 2 recruitment to CpG Islands genome wide. Mol Cell 55: 347–360

Roubille S, Escure T, Juillard F, Corpet A, Néplaz R, Binda O, Seurre C, Gonin M, Bloor S, Cohen C, et al (2024) The HUSH epigenetic repressor complex silences PML nuclear body-associated HSV-1 quiescent genomes. Proc Natl Acad Sci 121

Rowitch DH & Kriegstein AR (2010) Developmental genetics of vertebrate glial-cell specification. Nature 468: 214–222

Roy A & Ghosh A (2024) Epigenetic Restriction Factors (eRFs) in Virus Infection. Viruses 16: 183

Schuettengruber B, Bourbon H-M, Di Croce L & Cavalli G (2017) Genome Regulation by Polycomb and Trithorax: 70 Years and Counting. Cell 171: 34–57

Seczynska M, Bloor S, Cuesta SM & Lehner PJ (2022) Genome surveillance by HUSH-mediated silencing of intronless mobile elements. Nature 601: 440–445

Seczynska M & Lehner PJ (2023) The sound of silence: mechanisms and implications of HUSH complex function. Trends Genet 39: 251–267

Stovner EB & Sætrom P (2019) Epic2 efficiently finds diffuse domains in ChIP-seq data. Bioinformatics 35: 4392–4393

Stürzl M, Gaus D, Dirks WG, Ganem D & Jochmann R (2013) Kaposi’s sarcoma-derived cell line SLK is not of endothelial origin, but is a contaminant from a known renal carcinoma cell line. Int J Cancer 132: 1954–1958

Sun R, Tan X, Wang X, Wang X, Yang L, Robertson ES & Lan K (2017) Epigenetic Landscape of Kaposi’s Sarcoma-Associated Herpesvirus Genome in Classic Kaposi’s Sarcoma Tissues. PLoS Pathog 13: 1–21

Tischer BK, Smith GA & Osterrieder N (2010) En Passant Mutagenesis: A Two Step Markerless Red Recombination System. In Methods in Molecular Biology, Braman J (ed) pp 421–430. Totowa, NJ: Humana Press

Toth Z, Brulois K, Lee H-R, Izumiya Y, Tepper C, Kung H-J & Jung JU (2013) Biphasic Euchromatin-to-Heterochromatin Transition on the KSHV Genome Following De Novo Infection. PLoS Pathog 9: e1003813

Toth Z, Maglinte DT, Lee SH, Lee H-R, Wong L-Y, Brulois KF, Lee S, Buckley JD, Laird PW, Marquez VE, et al (2010) Epigenetic analysis of KSHV latent and lytic genomes. PLoS Pathog 6: e1001013

Uppal T, Banerjee S, Sun Z, Verma S & Robertson E (2014) KSHV LANA—The Master Regulator of KSHV Latency. Viruses 6: 4961–4998

Wachter E, Quante T, Merusi C, Arczewska A, Stewart F, Webb S & Bird A (2014) Synthetic CpG islands reveal DNA sequence determinants of chromatin structure. Elife 3: e03397

Wassef M, Luscan A, Aflaki S, Zielinski D, Jansen PWTC, Baymaz HI, Battistella A, Kersouani C, Servant N, Wallace MR, et al (2019) EZH1/2 function mostly within canonical PRC2 and exhibit proliferation-dependent redundancy that shapes mutational signatures in cancer. Proc Natl Acad Sci 116: 6075–6080

Weber K, Bartsch U, Stocking C & Fehse B (2008) A Multicolor Panel of Novel Lentiviral “Gene Ontology” (LeGO) Vectors for Functional Gene Analysis. Mol Ther 16: 698–706

Wu X, Johansen JV & Helin K (2013) Fbxl10/Kdm2b Recruits Polycomb Repressive Complex 1 to CpG Islands and Regulates H2A Ubiquitylation. Mol Cell 49: 1134–1146

Zhang Y, Liu T, Meyer CA, Eeckhoute J, Johnson DS, Bernstein BE, Nussbaum C, Myers RM, Brown M, Li W, et al (2008) Model-based analysis of ChIP-Seq (MACS). Genome Biol 9: R137

Zhu Y, Wang GZ, Cingöz O & Goff SP (2018) NP220 mediates silencing of unintegrated retroviral DNA. Nature 564: 278–282

